# Microglial 25-hydroxycholesterol mediates neuroinflammation and neurodegeneration in a tauopathy mouse model

**DOI:** 10.1101/2023.09.08.556884

**Authors:** Danira Toral-Rios, Justin M Long, Jason D. Ulrich, Jinsheng Yu, Michael R. Strickland, Xianlin Han, David M Holtzman, Anil G Cashikar, Steven M Paul

**Affiliations:** Department of Psychiatry, Washington University School of Medicine, St Louis, MO 63110; Department of Neurology, Washington University School of Medicine, St Louis, MO 63110; Hope Center for Neurological Disorders, Washington University School of Medicine, St Louis, MO 63110; Knight Alzheimer Disease Research Center, Washington University School of Medicine, St Louis, MO 63110; Genome Technology Access Center at the McDonnell Genome Institute and Department of Genetics, Washington University School of Medicine, St Louis, MO 63110; Barshop Institute for Longevity and Aging Studies, Department of Medicine, University of Texas Health Science Center, San Antonio, TX 78229; Taylor Family Institute for Innovative Psychiatric Research, Washington University School of Medicine, St Louis, MO 63110

## Abstract

Alzheimer’s disease (AD) is characterized by amyloid plaques and neurofibrillary tangles in addition to neuroinflammation and changes in brain lipid metabolism. Recent findings have demonstrated that microglia are key drivers of neurodegeneration in tauopathy mouse models. A subset of microglia referred to as disease-associated microglia (DAM) display gene signatures signifying changes in proinflammatory signaling and lipid metabolism in mouse models of amyloid and tau pathology. *Ch25h* is a DAM gene encoding cholesterol 25-hydroxylase that produces 25-hydroxycholesterol (25HC), a known modulator of inflammation as well as lipid metabolism. However, whether Ch25h influences tau-mediated neuroinflammation and neurodegeneration is unknown. Here, we show that in the absence of Ch25h and the resultant reduction in 25HC there is strikingly reduced age-dependent neurodegeneration and neuroinflammation in the hippocampus and entorhinal/piriform cortex of PS19 mice, which express the P301S mutant human tau transgene. Transcriptomic analyses of bulk hippocampal tissue and single nuclei revealed that Ch25h deficiency in PS19 mice strongly suppressed proinflammatory cytokine and chemokine signaling in microglia and restored sterol synthesis. Our results suggest a key role for Ch25h/25HC in potentiating proinflammatory signaling to promote tau-mediated neurodegeneration. Ch25h may represent a novel therapeutic target for primary tauopathies, AD, and other neuroinflammatory diseases.

## INTRODUCTION

Alzheimer’s disease (AD) is a progressive neurodegenerative disease histopathologically characterized by extracellular amyloid plaques containing aggregated forms of amyloid beta (Aβ)-peptides and intracellular neurofibrillary tangles composed mainly of hyperphosphorylated, aggregated tau. The onset and progression of tau pathology correlates with synaptic dysfunction and neuronal loss, leading to region-specific brain atrophy and cognitive impairment (1). Neuroinflammation is a prominent neuropathological feature of AD and considerable genetic and experimental evidence have implicated microglia, the brain’s resident immune cell, as major drivers of innate immunity-induced neurodegeneration (2). More than half of the reported genetic risk factors of AD are related to microglia-mediated immune response and cholesterol metabolism (3). Two of the major genetic risk factors for late onset AD, apolipoprotein E (*APOE*), and the triggering receptor expressed on myeloid cells 2 (*TREM2*), are both highly expressed in a unique subset of microglia termed disease-associated microglia (DAM) (4). A two-stage activation of microglial phenotype, switching from a homeostatic state to DAM has been shown to be *Apoe*- and *Trem2*-dependent (4–6). Deficiency of *Apoe* or *Trem2* attenuates tau-dependent neuroinflammation and neurodegeneration in mice expressing human P301S mutant tau (PS19 mice) (7–9). In addition, pharmacological depletion of microglia in the same transgenic mice decreased the age-dependent progression of tau pathology, indicating that tau-dependent neurodegeneration may occur primarily as a result of microglial activation leading to a state of chronic neuroinflammation and subsequent neuronal cell death (10). Data also suggest that activated microglia may also secrete one or more factors leading to neurodegeneration (2, 11).

Identification of the microglial factors that drive neuroinflammation-induced neurodegeneration is thus critically important and could lead to novel therapeutic strategies for treating or preventing AD and other tauopathies. In the current work, we focus on cholesterol 25-hydroxylase (Ch25h), which is upregulated in DAM only when both Apoe and Trem2 are present (5, 12). Ch25h converts cholesterol to 25-hydroxycholesterol (25-HC) (13). Mice lacking *Ch25h* appear to have a normal phenotype, show no changes in serum cytokine levels (14) and have normal cholesterol metabolism (15). Ch25h is primarily expressed in myeloid cells under inflammatory conditions and has been shown to have several important immune-related functions involving both innate and adaptive immunity (16). *Ch25h* is an important interferon (IFN)-inducible gene and 25-HC is believed to mediate the anti-viral properties of IFN in response to infection with a variety of viral pathogens (17, 18). 25-HC has also been reported to have both anti-(15, 19) and pro-inflammatory (20, 21) actions depending on the model system and context. In the central nervous system (CNS), Ch25h is upregulated in various mouse models of neuroinflammation (reviewed in [The Myeloid Landscape 2](http://research-pub.gene.com/BrainMyeloidLandscape/BrainMyeloidLandscape2/#Mouse-gene/Mouse-gene/12642/geneReport.html) database reported in (22). *Ch25h* was reported to be upregulated in patient-derived cells of X-linked adrenoleukodystrophy and administration of 25-HC into the corpus callosum of mice induced potent pro-inflammatory activation of microglia, NLRP3 inflammasome-mediated neuroinflammation, interleukin 1 beta (IL-1β) secretion and cell death (23). We previously reported that CH25H is upregulated in AD brain as well as mouse models of amyloid and tau pathology (24). We further demonstrated that mouse primary microglia treated with the TLR4 agonist lipopolysaccharide (LPS) produce 25-HC that markedly augment IL-1β production in an ApoE isoform-dependent manner (E4>>E3>E2) (24). Based on these findings, we hypothesized that 25-HC acts as a proinflammatory mediator and thus may contribute to neuroinflammation and neurodegeneration characteristic of various tauopathies including AD. To test this hypothesis, we generated Ch25h-deficient PS19 mice and show that Ch25h deficiency markedly attenuates neurodegeneration as well as significantly reduces the associated microgliosis and astrogliosis. Further, transcriptomic studies revealed marked suppression of inflammatory pathways and restored sterol biosynthesis in PS19 mice lacking Ch25h suggesting a major role for 25-HC in mediating changes in lipid metabolism, neuroinflammation and neurodegeneration. As a non-essential DAM gene, Ch25h may also represent a novel therapeutic target for AD and related tauopathies.

## RESULTS

### Expression of Ch25h in the brains of AD patients and PS19 mice

We previously reported increased expression of cholesterol-25-hydroxylase (CH25H) mRNA in the frontal cortex of AD patients as well as in mouse models of amyloid deposition and tau-mediated neurodegeneration (24). Although single cell transcriptomic analysis has identified CH25H as a marker of disease-associated microglia (DAM) in AD patients and transgenic mouse models bearing amyloid pathology (4, 5, 12) or tau pathology (22), cell-specific expression of the Ch25h protein has not been demonstrated in AD brain. Hence, we examined the localization of the Ch25h protein in relation to markers for microglia and astrocytes by immunostaining in the frontal cortex of AD brain tissues at Braak stages I-II and VI (Figure 1A, B and C, Supplementary Table I). Immunostaining with AT8 antibody, which recognizes phosphorylated tau (p-tau) on serine 202 and threonine 205 (Ser202/Thr205), showed that Braak stage VI samples had higher levels of p-tau relative to Braak I-II stage samples as expected (Figure S1A). CH25H expression was expected at both Braak stages since amyloid plaques are present in all the analyzed samples (Supplementary Table I) and amyloid triggers glia activation. However, the total volume of CH25H immunoreactivity was significantly increased (p<0.01) in Braak VI stage brain sections (33338 μm^3^, 95% CI 24997-41678) compared with Braak I-II (15047 μm^3^, 95% CI 3117-26977). To identify CH25H-cell specific expression a colocalization volume was created from CH25H and IBA1 immunoreactivity, and CH25H and GFAP immunoreactivity. About 90% of Ch25h immunoreactivity was observed within Iba1-positive microglia (Figure 1A, 1B and 1D) and less than 1% in GFAP-positive astrocytes (Figure 1A and 1E). Increased CH25H volume observed in microglia in late stages of AD suggest that tau pathology and associated neurodegeneration can induce Ch25h expression. Related to this, we previously identified an increased expression of Ch25h in the cortex of 9.5-month old PS19 mice (24), a tauopathy mouse model that overexpresses human tau containing the P301S mutation that causes a familial form of frontotemporal dementia (25). Here we conducted a *Ch25h* RNA in-situ hybridization assay (RNAscope, Figure 1G) in combination with immunostaining for Iba1 in the hippocampus of 9.5-month-old female PS19 (T) mouse and aged-match wild type (WT). The specificity of the Ch25h probe for RNAscope^®^ was validated by comparing the *Ch25h* mRNA signal in brain sections from WT and mice deficient for Ch25h (CKO). A low level of background reactivity was observed in the CKO mouse tissue (Figure S1B), similar to that observed using the negative control assay (for *Bacillus subtilis* dihydrodipicolinate reductase gene) RNAscope^®^ probe in CKO mouse tissue (Figure S1C), and was probably related to autofluorescence displayed in aged-mouse brain tissue. However, the RNAscope method allowed us to detect higher levels of Ch25h expressed mainly in microglia in PS19 mice compared to wild type mice (Figure 1G), similar to that observed in human brain tissue.

**Figure 1:**
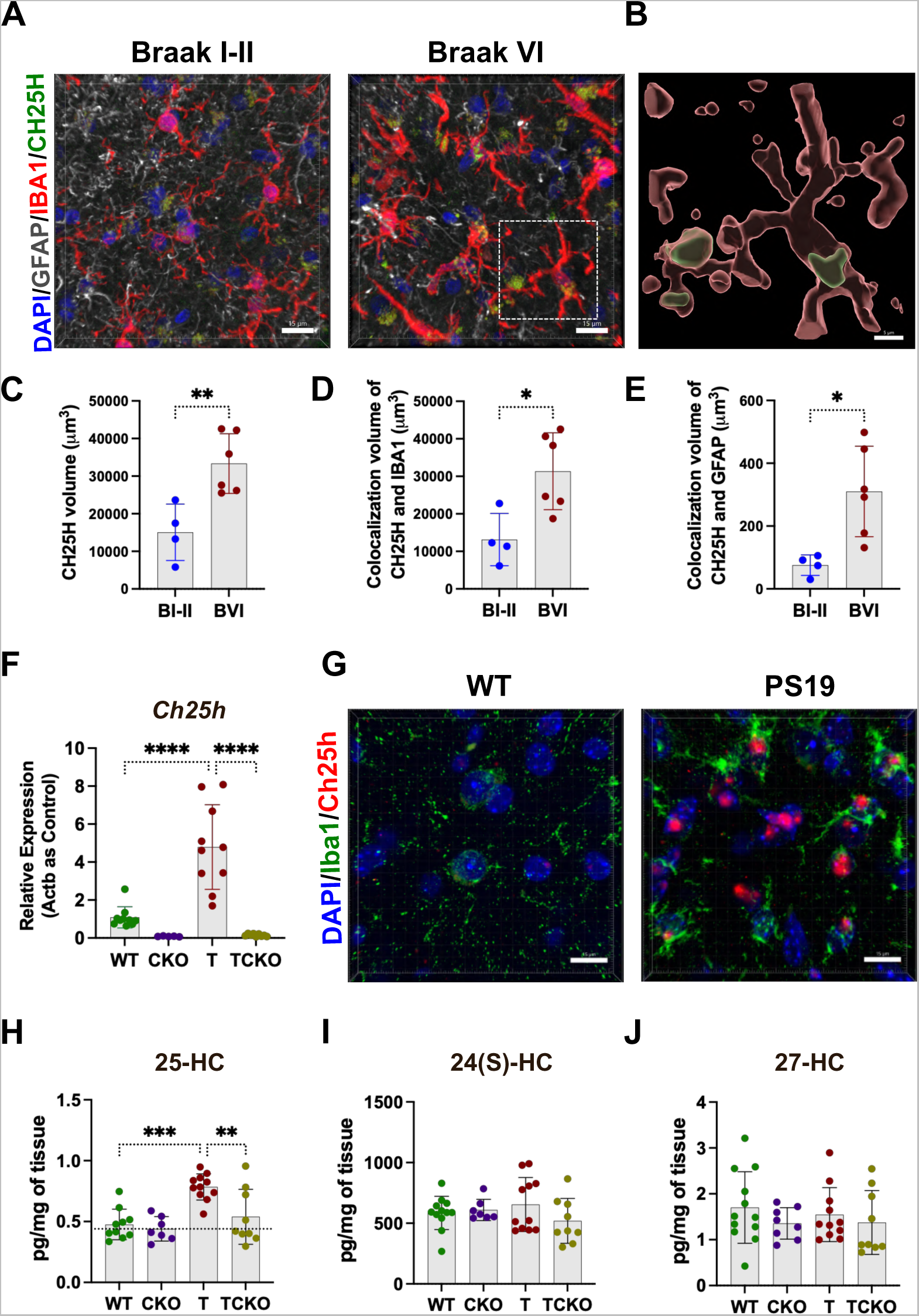
Ch25h is expressed mainly in microglia in brain tissue from AD patients and PS19 mice. A) Representative images of Braak I-II and Braak VI stage frontal cortex sections stained with GFAP, IBA1 and CH25H (scale bar 15 µm). A magnified surface 3D view (B) of a single microglial cell (Braak VI) and CH25H surface is shown (scale bar 5 µm). Total volume of CH25H immunoreactivity (C), colocalization volume of CH25H and IBA1-positive microglia (D), and colocalization volume of CH25H and GFAP-positive astrocytes (E) were quantified. Ch25h mRNA relative levels (F) were measured by qPCR in the hippocampus of WT(n=10), *Ch25h* knockout mice (CKO, n=5), T (n=10) and PS19 mice lacking Ch25h (TCKO, n=10) mice. Gene expression was normalized with β-actin (Actb). Representative images (scale bar 15 µm) from the hippocampus of 9.5-month old female wild type (WT) and PS19 (T) mouse stained with a *Ch25h* RNAscope assay (G), DAPI and Iba1. Levels of 25-hydroxycholesterol (H), 24-hydroxycholesterol (I) and 27-hydroxycholesterol (J) were quantified in the hippocampus of 9.5-month female WT (n=11), CKO (n=7), T (n=11) and TCKO (n=9) mice. Data expressed as mean ± SD, One-way ANOVA with Tukey’s post hoc test (two-sided) was used for all statistical analysis *p<0.05, **p<0.01, ****p<0.0001.

To investigate whether Ch25h affects tau-mediated neuroinflammation and neurodegeneration, we generated and aged PS19 female mice lacking Ch25h (TCKO), PS19 mice (T), Ch25h knock out (CKO), and wild-type (WT) mice. We quantified relative Ch25h mRNA levels by qPCR in the hippocampus of tau transgenic and non-transgenic female mice at 6 (Figure S1D) and 9.5 months (Figure 1F) of age. Although it has been reported that at 6 months of age only synaptic dysfunction (but no neurodegeneration), is observed due to tau accumulation in neurons accompanied by an increase in gliosis (25), no significant changes in Ch25h mRNA levels were detected in T mice relative to aged-matched WT mice (Fig S1D). By 9.5 months of age, a stage when PS19 mice show marked neuroinflammation and neurodegeneration, we identified a 3-4 fold increase in Ch25h expression in T mice compared to controls (p<0.0001), as well as the absence of Ch25h expression in knockout mice. Several transcriptomic studies have shown that Ch25h is primarily expressed by microglia in the CNS (22). Our previous studies with primary microglia isolated from wildtype and Ch25h-deficient mice show that only wildtype microglia produce and secrete high levels of 25-HC upon stimulation with an inflammatory challenge such as LPS (24). However, we cannot exclude the possibility that other cells such as astrocytes (26) and oligodendrocytes (23) can express Ch25h in the context of inflammation, or other disease-related pathology. However, our results suggest that Ch25h is mainly expressed in activated microglia in brains of AD and PS19 mice.

To confirm that changes in Ch25h expression resulted in an increase in 25-hydroxycholesterol (25-HC) synthesis, levels of 25-HC were quantified by LC/MS in the hippocampus of mice at the same ages (Figure 1H, S1E). Consistent with expression of Ch25h, a significant increase (p<0.0001) in the amount of 25HC was observed only in PS19 mice by 9.5 months of age compared with WT mice (Figure 1H). The low levels of 25-HC observed in the brain tissue from all mice may be produced by other cytochrome P450 enzymes (CYP3A4, CYP27A1, and CYP46A1) or via non-enzymatic oxidation (27). Levels of two oxysterol isomers of 25-HC, namely, 24S-hydroxycholesterol (24S-HC) and 27-hydroxycholesterol (27-HC), were also measured at 9.5-month (Fig 1I, 1J) and 6-month-old mice (Fig S1F, S1G). Although both 24S-HC and 27-HC are produced in the CNS, they were unaffected by tau pathology. Taken together, our results indicate that Ch25h expression and 25-HC synthesis in microglia in the brains of P301S mouse increase after the onset of tau pathology and are associated with neuroinflammation and neurodegeneration.

### Ch25h deficiency blocks tau-mediated neurodegeneration in PS19 mice

PS19 mice develop a profound loss of neurons in the hippocampus and entorhinal cortex by nine months of age, accompanied by tau hyperphosphorylation and aggregation, neurofibrillary tangle formation and gliosis (25). To determine whether Ch25h influences tau-mediated neurodegeneration in PS19 mice, we measured regional brain volumes in WT, CKO, T and TCKO mice (Figure 2A). As expected, T mice developed marked brain atrophy primarily of the hippocampus and entorhinal/piriform cortex compared with WT and CKO mice (Figure 2B and 2C), while there was a trend for increased ventricular volume that was not statistically significant (p=0.226) (Figure 2D).

**Figure 2:**
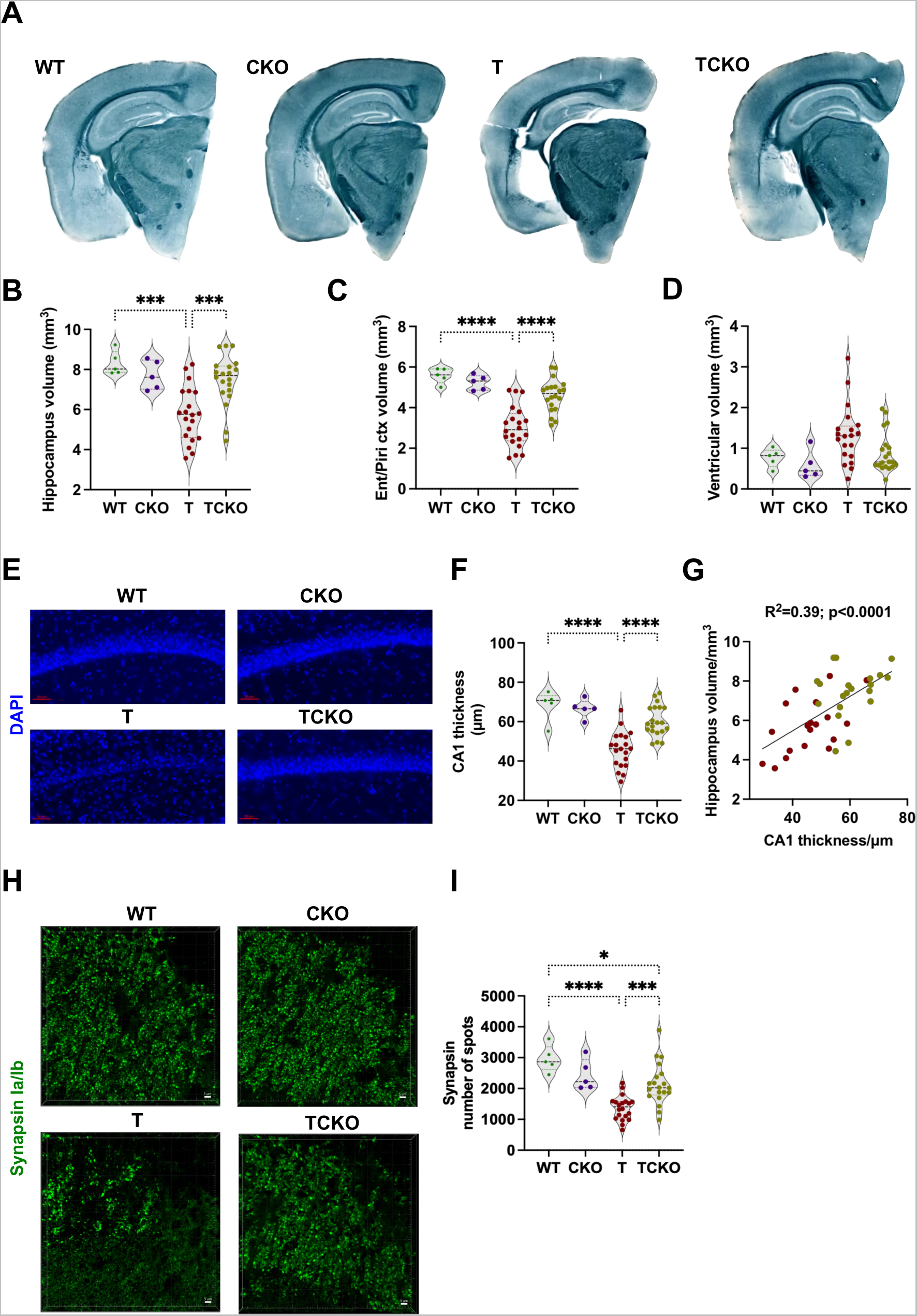
Ch25h deficiency reduces brain atrophy and neuronal loss in PS19 mice. A) Representative images of 9.5-month old wild type (WT), Ch25h KO (CKO), PS19 (T) and PS19/Ch25h KO (TCKO) mouse brain sections stained with Sudan black (WT: n=5, CKO: n=5, T: n=20, TCKO: n=20). Brain volume analysis of the hippocampus (B), piriform/entorhinal cortex (C) and posterior lateral ventricle (D). Representative images (E) of mouse brain sections of the CA1 region stained with DAPI (scale bar 50 µm). The thickness of the pyramidal cell layer is shown in (F). Pearson’s correlation analysis (two sided) between CA1 cell layer thickness with the hippocampal volume is shown (G) for tau mice - T (red) and TCKO (asparagus). Representative images of mouse brain sections stained with synapsin Ia/Ib antibody in the CA3 region (stratum lucidum) of the hippocampus (H). Quantification of the percentage of area covered by synapsin immunoreactivity (I). Data expressed as mean ± SD, One-way ANOVA with Tukey’s post hoc test (two-sided) was used for all statistical analysis *p<0.05, ***p<0.001, ****p<0.0001.

A comparison of hippocampal volumes in WT (8.301 mm^3^, 95% CI 7.559-9.043), T (5.605 mm^3^, 95% CI 4.971-6.239) and TCKO (7.466 mm^3^, 95% CI 6.880-8.052) mice indicates that hippocampal atrophy in TCKO mice was reduced (p<0.001) compared to T mice by approximately 70% (percentage was calculated with the formula – [(TCKO-T)/(WT-T)] X 100). Similar calculations in the entorhinal cortex reveal a reduction in brain atrophy in TCKO mice relative to T mice (p<0.001) by 62%.

Next, we assessed neuronal loss by measuring the thickness of the pyramidal cell layer in the hippocampal CA1 region (Figure 2E), the granule cell layer (Figure S2A) of the dentate gyrus (DG), as well of the pyramidal cell layer of the piriform cortex (Figure S2D). Similar to what we observed for hippocampal and cortical atrophy, there was a significant loss of neurons in the granule (Figure S2B) and pyramidal cell layers (Figure 2F and S2E) in T mice related to the control (WT) group. Neuronal loss in TCKO mice was markedly reduced (≥80%) and was similar to the non-transgenic control group. The comparison of CA1 pyramidal layer thickness in WT (68.38 μm, 95% CI 58.84-77.92), T (45.76 μm, 95% CI 41.48-50.04) and TCKO (60.33 μm, 95% CI 56.65-64.00) mice indicates that neuronal loss in TCKO mice relative to T mice was reduced (p<0.001) by approximately 64% (Figure 2F). The percent reduction in neuronal loss observed for DG granular layer and the pyramidal layer of the piriform cortex were 55% (Figure S2B) and 80%, respectively (Figure S2E). Changes in CA1 layer thickness correlated highly with hippocampal volumes (Figure 2G; correlation coefficient = 0.39, p<0.0001), as did DG layer thickness (Figure S2C; correlation coefficient = 0.37, p<0.0001), and the pyramidal layer of the piriform cortex thickness correlated with piriform/entorhinal cortex volume (Figure S2F; correlation coefficient = 0.58, p<0.0001).

Synaptic loss was evaluated in the stratum lucidum of hippocampal CA3 region by immunostaining for the presynaptic protein synapsin (Figure 2H). A marked reduction of synapsin immunoreactivity was found in T and TCKO mice compared with WT mice (Figure 2I). However, synapsin loss was significantly attenuated in TCKO mice relative to T mice (p<0.001). Progression of tau pathology has been correlated with memory dysfunction and cognitive impairments in AD (28, 29). In PS19 mice, early intracellular aggregation of tau can impair long-term potentiation (LTP), a mechanism related to synaptic plasticity and consolidation of memory (25). Soluble factors secreted by activated microglia under conditions of chronic inflammation have been thought to mediate aberrant synaptic phagocytosis inducing synapse loss and cognitive impairment (30, 31). Related to this, we recently demonstrated that 25-HC disrupts synaptic plasticity and learning in mice treated with LPS (32) suggesting the possibility that 25-HC may trigger tau-induced impairment of synaptic plasticity in addition to neuronal loss. Nonetheless, the striking reduction in brain atrophy, neuronal and synaptic loss in TCKO mice underscores an important role for Ch25h and 25-HC in the development of tau-mediated neurodegeneration.

### Ch25h deficiency reduces tau phosphorylation and aggregation in PS19 mice but doesn’t affect tau seeding and spreading

To determine whether the reduced neurodegeneration in TCKO mice was accompanied by a reduction in tau phosphorylation, we stained tissue sections with the AT8 antibody. As expected, p-tau was undetectable in WT and CKO mice and detected only in T and TCKO, which overexpressed human-mutant tau (Figure 3A). Quantification of the percentage of area covered by AT8 immunoreactivity revealed a significant decrease in AT8-positive p-tau in PS19 mice lacking Ch25h in both the hippocampus and entorhinal/piriform cortex (Figure 3C, 3D). The comparison of the percentage of AT8 immunoreactive area in the hippocampus of WT (0.002%, 95% CI 0.001-0.004), T (19.59%, 95% CI 14.0-25.17) and TCKO (7.97%, 95% CI 5.404-10.52) mice indicates that AT8 immunoreactivity in TCKO mice was reduced (p<0.001) compared to T mice by approximately 60%. The AT8 immunoreactivity in the entorhinal/piriform cortex was reduced in TCKO mice (p<0.05) in comparison with T mice by 45%.

**Figure 3:**
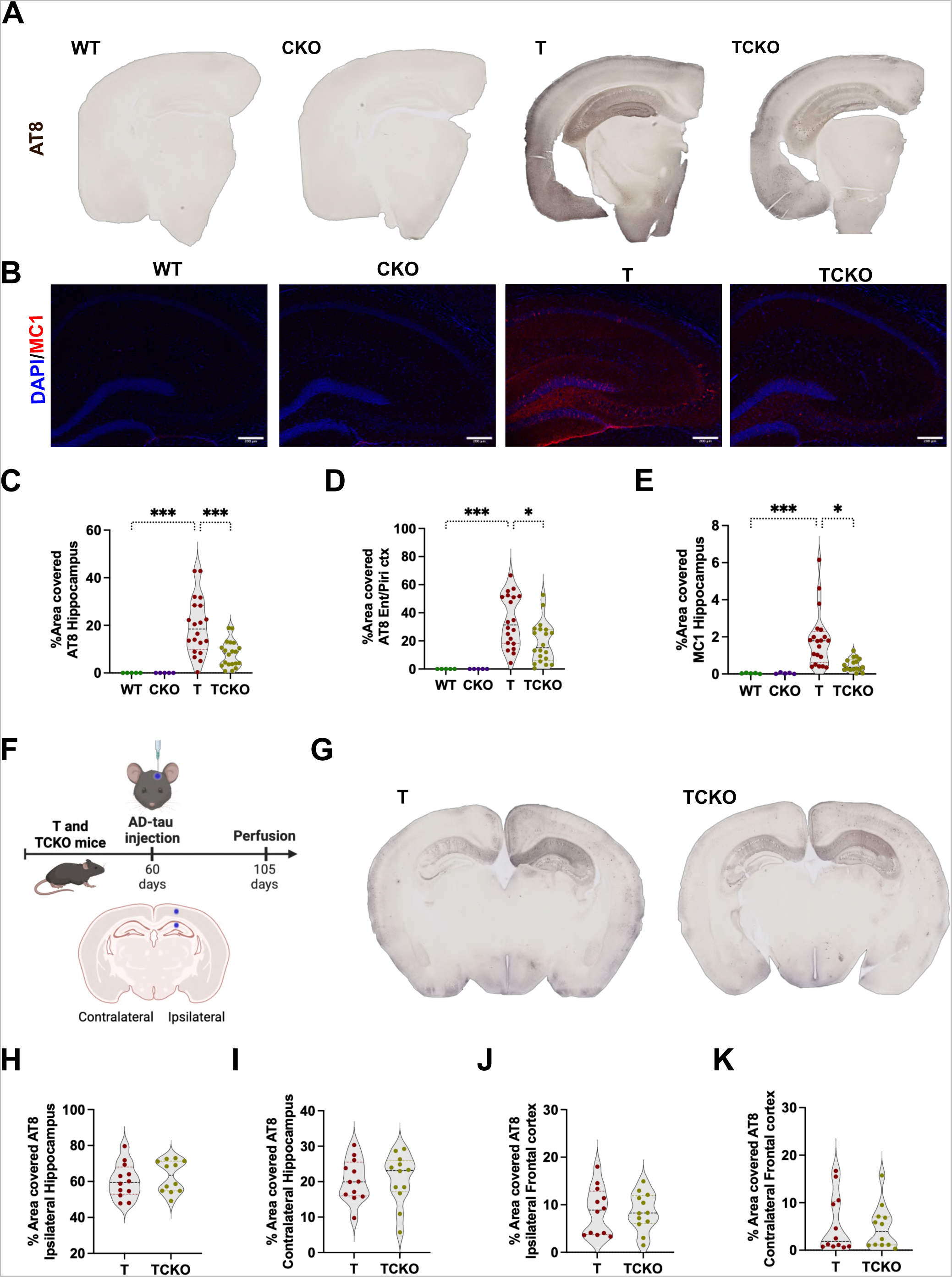
Ch25h deficiency reduces p-tau (Ser202/Thr205) immunoreactivity and tau aggregation in PS19 mice. A) Representative images of 9.5-month old wild type (WT), Ch25h KO (CKO), PS19 (T) and PS19/Ch25h KO (TCKO) mouse brain sections stained with AT8 antibody (WT: n=5, CKO: n=5, T: n=20, TCKO: n=20). B) Representative images showing DAPI (blue) and MC1 immunoreactivity (red) in the hippocampus of 9.5-month mouse brain sections (scale bar 200 µm). Quantification of the percentage of area covered with the AT8 immunoreactivity in the hippocampus (C) and entorhinal/piriform cortex (D). Percentage of area covered by MC1 immunoreactivity was quantified in the hippocampus (E). Schematic of tau seeding and spreading model with intracerebral injections of AD-tau in T and TCKO mice (F). Representative images from mouse brain sections immunostained with AT8 antibody (G). Quantification of the percentage of area covered with AT8 immunoreactivity was performed in the ipsi and contralateral hippocampus (H, I), as well as ipsilateral and contralateral frontal cortex (J, K). Data expressed as mean ± SD, One-way ANOVA with Tukey’s post hoc test (two-sided) was used for AT8 statistical analysis and Kruskal-Wallis test for MC1. *p<0.05, ***p<0.001. Ent: Entorhinal cortex, Piri: Piriform cortex.

Hyperphosphorylation of tau induces conformational changes that promotes protein aggregation. The conformational state of tau was analyzed by immunostaining with the MC1 antibody, which recognizes aggregates in an early stage of AD (Figure 3B). The area covered by MC1 immunoreactivity (Figure 3E) was significantly higher (p<0.001) in T mice (1.85%, 95% CI 1.152-2.564) relative to WT mice (0.03%, 95% CI 0.005-0.06). A significant reduction (75%) of MC1 immunoreactivity was observed in TCKO mice (0.49%, 95% CI 0.310-0.672) compared to T mice. Our results suggest that Ch25h-deficiency reduces tau phosphorylation and aggregation.

A protective role for 25-HC in blocking tau entry into neurons has been reported in a recent in vitro study (33). To determine whether Ch25h-deficiency affects tau seeding and spreading we used the in vivo model described by Boluda et al. (34), a model for tau seeding and spreading following intracerebral injection of AD brain extract enriched in pathological tau (AD-tau) in 2-month old T and TCKO mice (Figure 3F). This early age is prior to the onset of mutant tau transgene-induced pathology. Tau phosphorylation was evaluated by AT8 immunostaining in brain sections (Figure 3G). AD-tau injection induced abundant tau phosphorylation mainly in the ipsilateral hippocampus, where almost 60% of area covered by AT8 immunoreactivity was detected and was similar in T and TCKO mice (Figure 3H). Only 8% of the ipsilateral frontal cortex area was covered by AT8 immunoreactivity and no significant changes between T and TCKO mice were observed (Figure 3J). Tau phosphorylation was detected in the contralateral hippocampus and frontal cortex (10% and 5% of the area covered by AT8 immunoreactivity respectively) equally in T and TCKO mice (Figure 3I, 3K). Our results indicate that under our in vivo assay conditions Ch25h-deficiency does not alter tau seeding and spreading.

### Reduced gliosis in Ch25h deficient PS19 mice

By 9.5 months of age, PS19 mice manifest a marked microgliosis and astrogliosis which have been associated with tau phosphorylation and neurodegeneration (7, 25). We have recently shown that 25HC acts as an amplifier of the microglial inflammatory response (24) and can also regulate lipid metabolism in astrocytes (35). We hypothesized that 25HC, produced by Ch25h-expressing microglia, exerts autocrine and paracrine effects contributing to tau-dependent neuroinflammation. To test this, we quantified markers of astrocyte and microglial activation, and conducted an analysis of glial morphology. Sections immunostained for GFAP were assessed for astrogliosis (Figure 4A). A marked increase of the area covered by the GFAP immunoreactivity was observed in T mice compared to WT and CKO mice in the hippocampus (p<0.001; Figure 4C) and entorhinal/piriform cortex (p<0.001; Figure 4D). Although, the percentage of area covered by GFAP in TCKO mice was greater than that of non-tau mice (WT and CKO), the differences were not significant. However, the area of GFAP immunoreactivity in TCKO brain tissue was significantly lower than that in T mice in the hippocampus (p<0.01; Figure 4C) as well as entorhinal/piriform cortex (p<0.01; Figure 4D). In the astrocyte morphology analysis (Figure 4B) we identified numerous reactive astrocytes in T mice, characterized by a significant reduction (p<0.00001) in length of the processes (Figure 4E) and the number of branches per cell (Figure 4F). Astrocytes from TCKO mice displayed a significant reduction in both parameters in comparison with WT mice, and the processes were more ramified than astrocytes in T mice (p<0.00001), which is consistent with the reduction in GFAP immunoreactivity.

**Figure 4:**
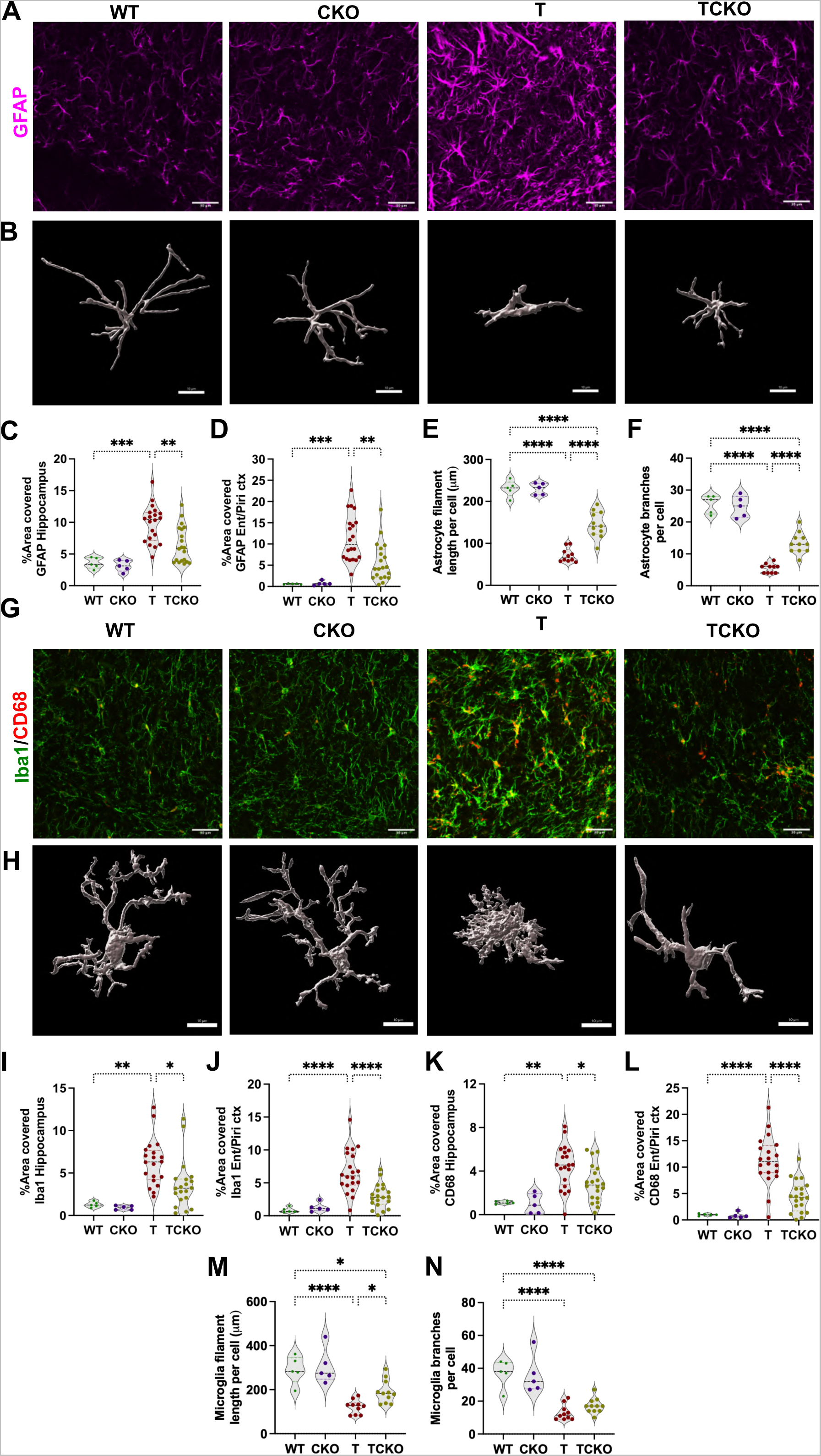
Ch25h deficiency decreases gliosis in PS19 mice. A) Representative images of sections stained with GFAP antibody (scale bar 30 µm) and (B) Imaris-3D reconstruction (Scale 10µm) from astrocyte morphology analysis to assess astrogliosis in 9.5-month old wild type (WT n=5), Ch25h KO (CKO n=5), PS19 (T n=20) and PS19/Ch25h KO (TCKO n=20) mice. Quantification of the percentage of area covered by GFAP immunoreactivity in the hippocampus (C) and (D) entorhinal/piriform cortex. For astrocyte morphology analysis, the filament length (E) and number of branches (F) per cell were quantified. G) Representative images of sections stained with Iba1 (green) and CD68 (red) to assess microgliosis (scale bar 30 µm). H) Imaris-3D reconstruction from a representative Iba+ microglia per group (scale bar 10 µm). Quantification of the percentage of area covered by Iba1 (I, J) and CD68 (K, L) immunoreactivity in the hippocampus (I, K) and entorhinal/piriform cortex (J, L). Quantification of microglial filament length per cell (M) and branches per cells (N) are shown. Data expressed as mean ± SD, One-way ANOVA with Tukey’s post hoc test (two-sided) was used for all statistical analysis *p<0.05, **p<0.01, ***p<0.001, ****p<0.0001. Ent: Entorhinal cortex, Piri: Piriform cortex.

To assess microgliosis, we immunostained sections for Iba1 together with cluster of differentiation 68 (CD68), a phagolysosome marker expressed by activated microglia (Figure 4G). The area of Iba1 and CD68 immunoreactivity were significantly elevated in T mice in agreement with previously published results (7). The percentage of surface area covered by Iba1 immunoreactivity was markedly reduced in TCKO mice in comparison with T mice in the hippocampus (p<0.05; Figure 4I) as well as in entorhinal/piriform cortex (p<0.0001; Figure 4J). The area covered by CD68 immunoreactivity was also significantly reduced in the hippocampus (p<0.05; Figure 4K) and entorhinal cortex (p<0.0001; Figure 4L) of TCKO mice. No significant differences were detected in Iba1 and CD68 immunoreactivity between TCKO and non-tau (WT and CKO) mice. Homeostatic microglia present a ramified morphology with longer processes, such as that observed in WT and CKO mice (Figure 4H, 4M, 4N), while microglia from T mice display a hypertrophic morphology characterized by a significant reduction in process length and number of branches per cell (p<0.0001). We found that microglia from TCKO mice also present a significant reduction in the number of branches per cell compared to the non-tau mice (Figure 4N), but the processes were in fact significantly longer (p<0.05) than microglia from T mice (Figure 4M), which suggests microglia in TCKO mice are less activated, possibly in an intermediate state of activation. Taken together, these results confirm that Ch25h deficiency results in significantly reduced astrocyte and microglial reactivity in PS19 transgenic mice.

### Ch25h deficiency reduces disease-associated microglia in PS19 mice

Single cell transcriptomic analysis carried out in AD human and AD mouse model brain tissue have identified unique signatures of gene expression in microglia in physical proximity to amyloid plaques and tau aggregates, termed disease-associated microglia (DAM) (4, 5, 12). This microglial subset is characterized by reduced expression of homeostatic genes (e.g. Tmem119, P2ry12, Cx3cr1) and increased expression of inflammatory genes (e.g. Clec7a, Apoe, Trem2) including Ch25h (36). Given the decrease in microglial reactivity observed in TCKO mice, we further investigated whether Ch25h deficiency favors a reduction in DAM in lieu of a more homeostatic phenotype in PS19 transgenic mice. To this end, we quantified Clec7a and Tmem119 immunoreactivity in the hippocampus and entorhinal cortex of T vs TCKO mice. The area covered by Clec7a immunoreactivity (Figure 5A) was substantially decreased (approximately 3-fold, p<0.05) in the hippocampus (Figure 5B) and entorhinal cortex of TCKO mice relative to T mice (Figure 5C). Clec7a mRNA levels in the hippocampus of TCKO mice were also reduced by 2-fold (p<0.01) in comparison with T mice (Figure 5D). No significant changes in Clec7a immunoreactivity were observed between TCKO and non-tau control (WT or CKO) mice. These results suggest that Ch25h and 25-HC may contribute to the transition of homeostatic to disease-associated microglia. We next quantified two additional DAM markers, ApoE (Figure S3A) and Trem2 (Figure S3D). As expected, ApoE immunoreactivity in the hippocampus increased markedly in T mice in comparison to control mice. Strikingly, ApoE immunoreactivity reverted to almost control levels in TCKO mice (Figure S3B). Given that ApoE is also normally expressed by astrocytes, sections were double-stained for Clec7a to specifically quantify the changes in ApoE immunoreactivity in activated microglia. Almost all the increased ApoE immunoreactivity in T mice localized to Clec7a-positive microglia (Figure S3C). Trem2 immunoreactivity (Figure S3E) was also found reduced in the hippocampus of TCKO mice relative to T mice (p<0.01). Likewise, Trem2 immunoreactivity in Clec7a-positive microglia was also significantly decreased in TCKO relative to T mice (p<0.01, Figure S3F).

**Figure 5:**
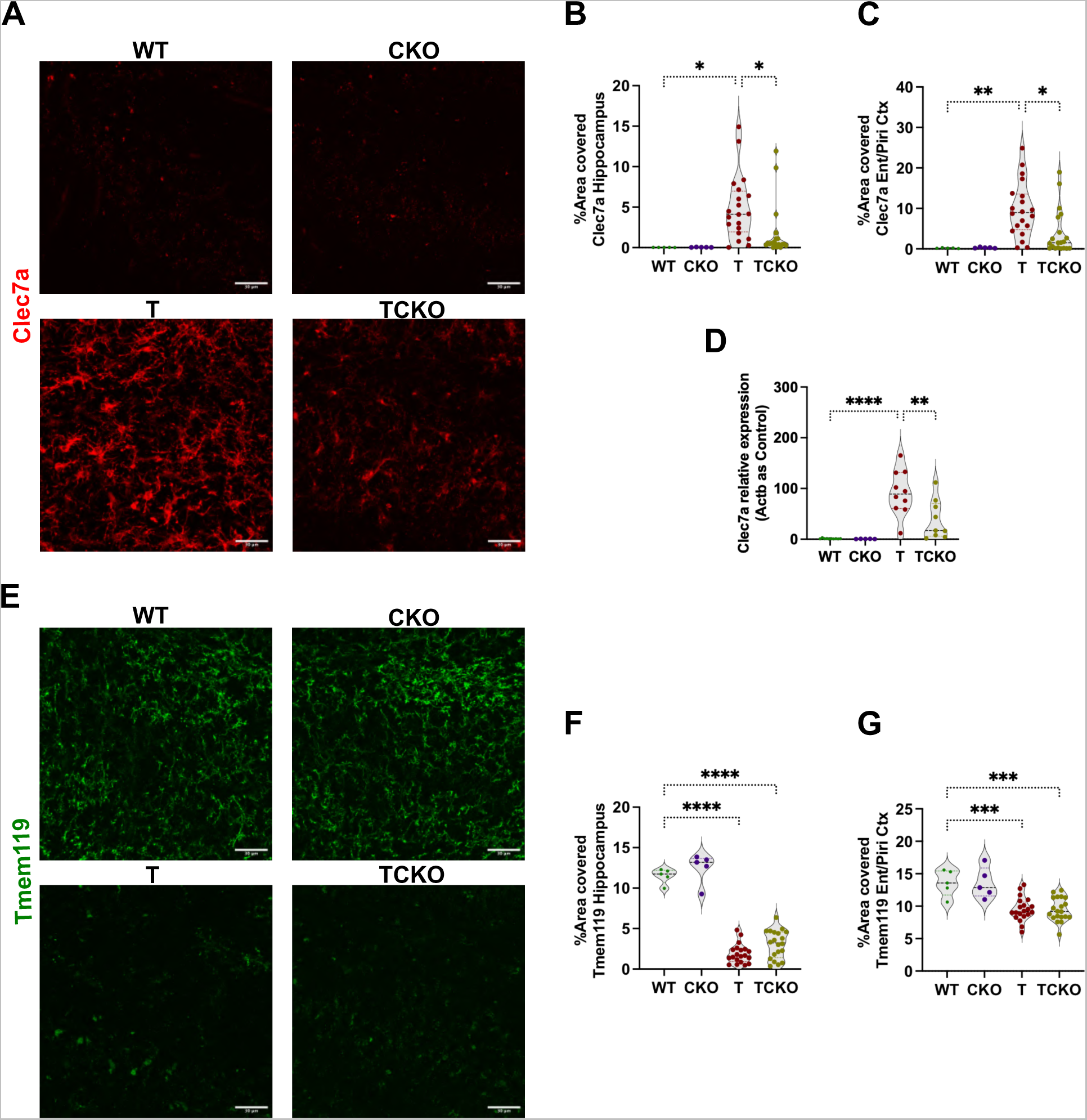
Ch25h deficiency decreases disease-associated microglia (DAM) in PS19 mice. A) Representative images of DAM microglia stained with Clec7a antibody in 9.5-month old wild type (WT, n=5), Ch25h KO (CKO, n=5), PS19 (T, n=20) and PS19/Ch25h KO (TCKO, n=20) mouse brain sections. Quantification of the percentage of area covered by Clec7a immunoreactivity in B) hippocampus and C) entorhinal/piriform cortex. D) Relative expression of Clec7a mRNA in the hippocampus was quantified by qPCR (WT: n=10, CKO: n=5, T: n=10, TCKO: n=10). Gene expression was normalized with ý-actin (Actb). (E) Representative images of homeostatic microglia immunostained for Tmem119. Quantification of the percentage of area covered by Tmem119 in the F) hippocampus and G) entorhinal/piriform cortex. Data expressed as mean ± SD. One-way ANOVA with Tukey’s post hoc test (two-sided) was used for all statistical analysis *p<0.05, **p<0.01, ***p<0.001, ****p<0.0001. Scale bar 30 µm. Ent: Entorhinal cortex, Piri: Piriform cortex.

Tau-mediated neuroinflammation also coincides with a marked decrease of homeostatic microglia. To test whether the decreased activation of microglia in TCKO is also associated with a reversal to a more homeostatic (resting) state, we stained tissue sections for Tmem119 (Figure 5E). In T mice, the area covered by Tmem119 immunoreactivity was significantly reduced relative to control (WT and CKO) mice. Interestingly, the area covered by Tmem119 immunoreactivity remained reduced in TCKO mice in the hippocampus (Figure 5F) and entorhinal/piriform cortex (Figure 5G). This correlates with our findings from the morphological analysis, suggesting that microglia in TCKO mice only partially reverted back to the homeostatic state. To confirm these observations, we immunostained for an additional homeostatic microglia marker, P2ry12 (Figure S3G) and total P2ry12 area was quantified (Figure S3H). Again, in T mice P2ry12 area was markedly decreased similar to Tmem119. In TCKO mice, P2ry12 area also remained low and did not revert to control (WT and CKO) levels. Overall, Ch25h deficiency in PS19 mice appears to strongly decrease progression of reactive microglia to a DAM state, but without fully reverting to a resting (homeostatic) microglial state.

### Ch25h deficiency results in transcriptomic changes involved in inflammation, cholesterol metabolism and trans-synaptic signaling in PS19 mice

We next profiled the transcriptomic changes associated with Ch25h deficiency by RNA-seq using RNA extracted from the hippocampus from WT, CKO, T and TCKO mice (n=4 mice per group). First, we compared the expression level of human transgenic *MAPT* gene in T and TCKO mice and found no significant differences indicating that the prevention of neuroinflammation and neurodegeneration in TCKO was not simply due to reduced expression of the tau transgene compared to PS19 mice. Differentially expressed genes (DEGs) filtered at p<0.05 and fold change (FC) >1.5 were analyzed for the following comparisons between groups: CKO vs WT, T vs WT, TCKO vs CKO and TCKO vs T. The volcano plots (Figure S4A) show differentially up- and down-regulated genes as well as expression of Ch25h. The numbers of DEGs overlapping in different comparisons are shown in the Venn diagram (Figure S4B, S4C and Supplementary table II). In Figure S4D, we present a heatmap including all 3918 DEGs between all 4 groups (gene lists are provided in Supplementary Table III). It is apparent that expression of the tau transgene results in increased expression of a large number of genes whereas only a smaller number of genes are downregulated in a tau-specific manner. Interestingly, the tau-specific changes in gene expression were mostly absent in TCKO mice. Some genes show increased or decreased expression in CKO and TCKO but not in WT and T mice suggesting that these DEGs are specifically changing due to the absence of Ch25h.

A comparison of DEGs between CKO and WT mice showed 295 upregulated genes and 305 downregulated genes (Figure S4A, S4B and S4D, Supplementary tables II, IV and V). However, there were no DEGs detected (up or down) when we corrected for false discovery rate (FDR, q<0.05), highlighting the fact that Ch25h is not highly expressed under non-inflammatory conditions and thus deleting it did not cause major transcriptional changes under normal conditions (Supplementary Table II).

In contrast, the presence of the tau transgene caused a much greater number of DEGs on a *Ch25h*+/+ background than on the *Ch25h*-/- background. In T versus WT comparison, 2477 genes were upregulated and 831 were downregulated (Figure S4B & S4C). However, only 770 genes were upregulated and 275 were downregulated in TCKO compared to CKO mice indicating that the tau transgene caused far fewer transcriptional changes in the absence of Ch25h. Further, comparing TCKO versus T mice there were 459 genes that were upregulated and 1634 genes that were downregulated relative to T mice (Figure S4C).

As expected, in T mice important DAM signature genes including *Apoe, Trem2, Cst7, Clec7a, Tyrobp, Axl, C1q, Csf1, Cstd and Cstb* were among those upregulated while in TCKO mice, a marked reduction of DAM gene expression was observed (Figure S5A, Supplementary Table III). The upregulated reactive astrocyte genes in T mice including *Gfap, Aqp4, Vim, Id3, Fabp7, Mt1, Mt2* were all decreased in TCKO mice (Figure S5B, Supplementary Table III). From this analysis it is apparent that the DAM and reactive astrocyte gene expression profiles in TCKO mice are similar to WT and CKO mice.

We next performed a pathway enrichment analysis using Metascape, a bioinformatics tool that incorporates a core set of default gene ontologies (37) (Figure 6, Supplementary Table IV and V). Neuroinflammatory genes were not the only subset of genes impacted by Ch25h loss in PS19 mice. Among the 831 DEGs that were downregulated in T vs. WT mice, genes belonging to the GO terms ‘trans-synaptic signaling’, ‘modulation of chemical synaptic transmission’ and ‘sterol biosynthesis’ were also observed (Figure 6E and 6G, Supplementary Table IV). Upregulation of these same pathways was observed in TCKO vs. T comparison (Figure 6B). Similar results were observed in the detailed heatmaps of individual DEGs belonging to the above pathways, where the gene expression levels were comparable to WT and CKO mice (Figure S4F and S4G). Analyses corresponding to the upregulated GO terms from the TCKO vs T dataset, namely ‘trans-synaptic signaling’ (Figure 6E and 6F) and ‘sterol biosynthesis’ (Figure 6G and 6H) revealed a single major subnetwork. In support of the observed DEGs in trans-synaptic signaling, we found that Ch25h deficiency prevented tau-mediated synaptic loss (Figure 2H and 2I).

**Figure 6:**
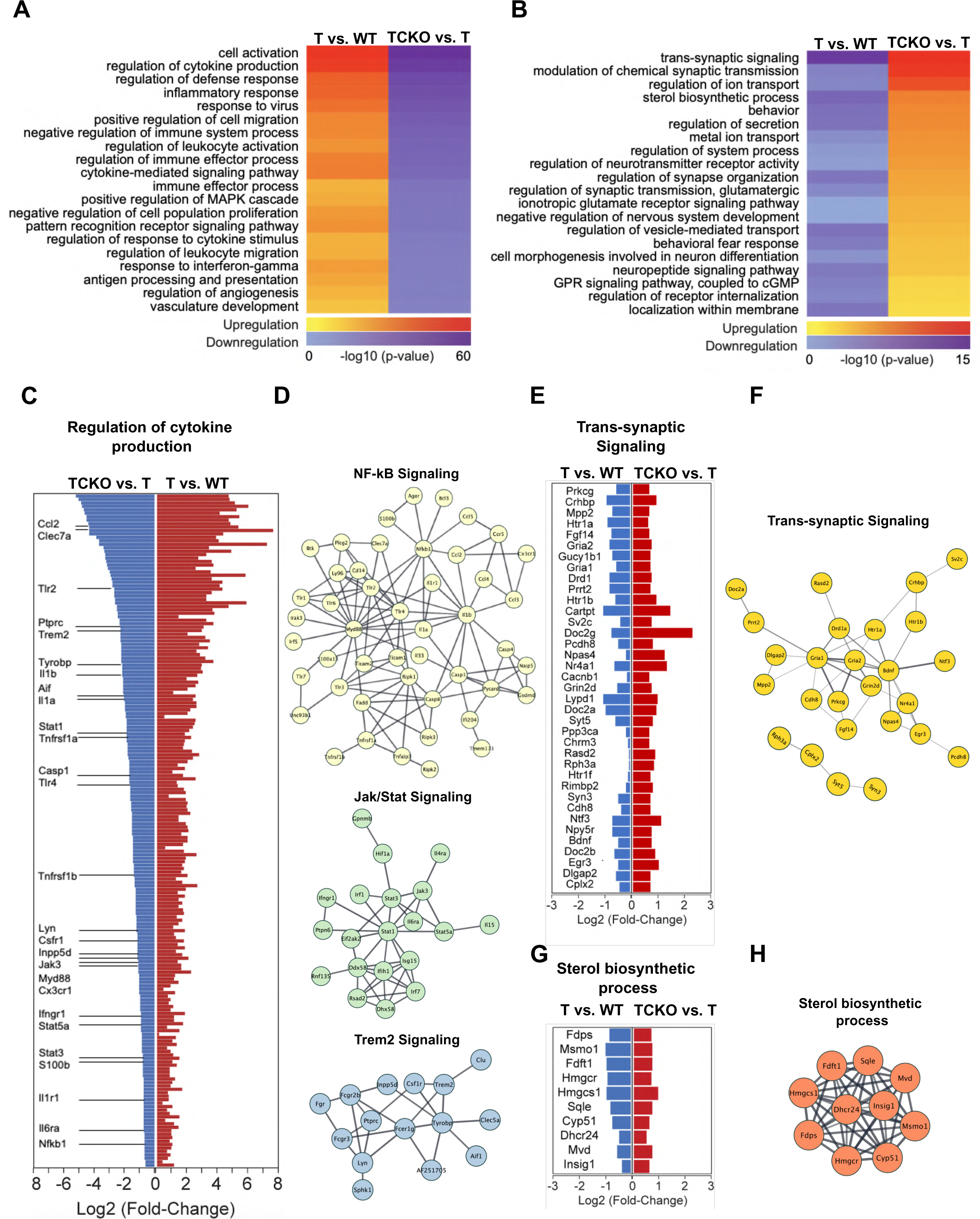
Ch25h deficiency results in downregulation of DEGs related to inflammatory processes and upregulation of DEGs related to trans-synaptic signaling and sterol biosynthesis in tau transgenic mice. Heatmaps of down (A) and up (B) regulated top gene ontology (GO) terms in T vs WT and TCKO vs T comparisons and arranged based on p-values from TCKO vs. T mice comparisons in a decreasing order. Shades of blue indicate downregulated GO terms and shades of red indicate upregulated GO terms. Plots of log2(fold change) in DEGs and interaction networks belonged to the GO terms ‘regulation of cytokine production’ (C and D), ‘trans-synaptic signaling’ (E and F) and ‘sterol biosynthetic process’ (G and H) between T vs. WT, and TCKO vs. T mice comparisons.

Gene ontology analysis of the upregulated DEGs in T versus WT mice, revealed pathways associated with inflammatory processes such as ‘regulation of cytokine production’, ‘leukocyte activation’, ‘regulation of defense response’ and ‘inflammatory response’ (Figure 6A, Supplementary Table IV). Analysis of the GO term ‘regulation of cytokine production’, showed genes that were upregulated in a tau-specific manner in comparison to non-tau wildtype mice and were all downregulated in Ch25h-deficient PS19 mice (Figures 6C and Supplementary Table V). Interestingly, the heatmap corresponding to the same GO term indicates that the transcriptome of TCKO hippocampi was more similar to WT and CKO mice than to the transcriptome of T mice (Figure S4E). Remarkably, almost complete restoration of the transcriptomic profile resulting from expression of the mutant tau transgene is observed in the absence of Ch25h despite the expression of the mutant tau transgene. Gene network analysis of the downregulated DEGs corresponding to ‘regulation of cytokine production’ from the TCKO vs T dataset, revealed pathways corresponding to ‘NF-kB signaling’, ‘Jak/Stat signaling’ and ‘Trem2 signaling’ (Figure 6D). This suggests an interplay between these proinflammatory pathways in tau-mediated neuroinflammation that is prevented by the absence of Ch25h.

We used the Transcriptional Regulatory Relationships Unraveled by Sentence-based Text mining (TRRUST) database to uncover the main transcription factors responsible for the differential expression of inflammatory pathway genes identified with bulk hippocampal tissue RNAseq, focusing on genes upregulated in T vs WT comparison and downregulated in TCKO vs T comparison. Transcription factors such as *Rela and Nfkb1 (both from NF-kB family), Trp53, Sp1, Jun, and Stat3* were among the top drivers of the transcriptional differences observed between TCKO and T mice (Figure 7A).

**Figure 7:**
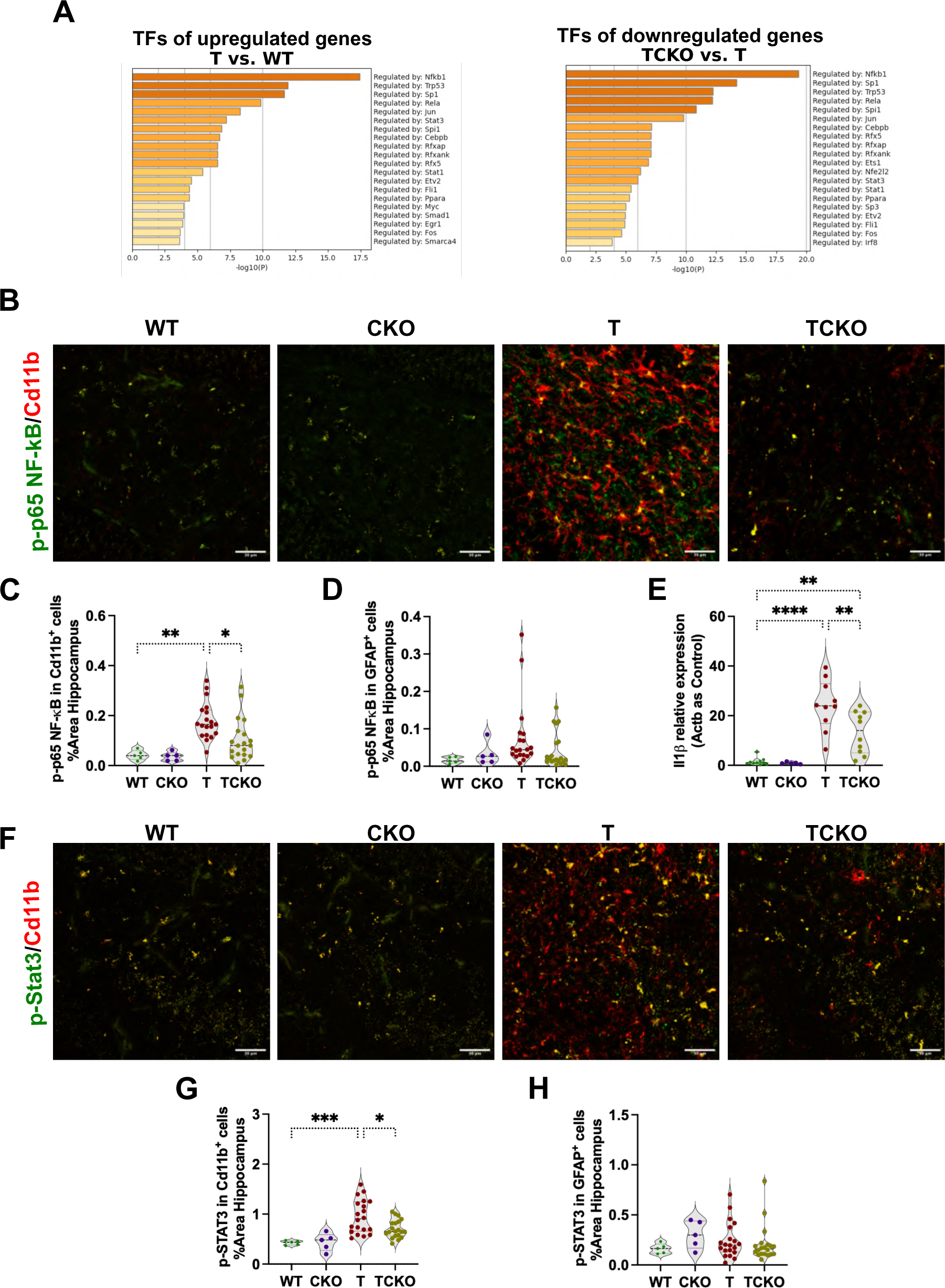
Ch25h deficiency in PS19 mice change the expression of key transcription regulators of the inflammatory response. A) TRRUST analysis of transcription factors (TFs) predicted to be responsible for the upregulated genes in T vs WT and downregulated genes in TCKO vs. T comparisons. B) Representative images from a double immunostaining for p-p65 NF-kB activated subunit (green) and the microglia marker Cd11b (red) in 9.5-month old wild type (WT, n=5), Ch25h KO (CKO, n=5), PS19 (T, n=19) and PS19/Ch25h KO (TCKO, n=19) mouse brain sections. Quantification of the percentage of area covered by p-p65 NF-kB immunoreactivity in Cd11b positive cells (C) and in GFAP positive cells (D) in the hippocampus. E) Relative expression IL-1ý (*Il1b*) mRNA levels were quantified in the hippocampus by qPCR (WT: n=10, CKO: n=5, T: n=10, TCKO: n=10). Gene expression was normalized with ý-actin (*Actb*). F) Representative images from a double immunostaining for p-STAT3 (green) and the microglia marker Cd11b (red) in 9.5-month old wild type (WT, n=5), Ch25h KO (CKO, n=5), PS19 (T, n=20) and PS19/Ch25h KO (TCKO, n=20) mouse brain sections. Quantification of the percentage of area covered by p-STAT3 immunoreactivity in Cd11b positive cells (G) and GFAP positive cells (H) in the hippocampus. Data expressed as mean ± SD. One-way ANOVA with Tukey’s post hoc test (two-sided) was used for all statistical analysis *p<0.05, **p<0.01, ***p<0.001, ****p<0.0001. Scale bar 30 µm.

### Single nuclei transcriptomics reveals the cellular origins of transcriptional changes

To better understand and corroborate cell-type-specific transcriptional changes associated with Ch25h-deficiency in neuroinflammation in PS19 mice, we performed single nuclei RNA-seq (snRNA-seq) using hippocampal tissue (5 mice pooled per group) from 6-month old T and TCKO mice, as well as 9.5-month-old WT, T and TCKO mice. Transcriptional data for 40,107 nuclei (Supplementary Table VII) was subjected to unsupervised clustering into 16 distinct clusters (Figures 8A, Supplementary Table VIII) based on the expression of cell-type-specific markers. The subclusters were categorized into excitatory neurons (exc 1,2,3,4,5,6,7), inhibitory neurons (inh 1,2,3), astrocytes (astro), microglia (micro), oligodendrocytes (oligo 1,2), oligodendrocyte precursor cells (opc) and choroid plexus epithelial cells (epi). Next, we examined the average expression of genes related to NF-kB signaling, JAK-STAT signaling and sterol biosynthesis for nuclei from specific cell-type composition (Figure 8B). Genes related to NF-kB and Jak/Stat pathways are highly expressed in microglia nuclei and genes related to sterol biosynthesis were highly expressed in astrocyte nuclei.

**Figure 8:**
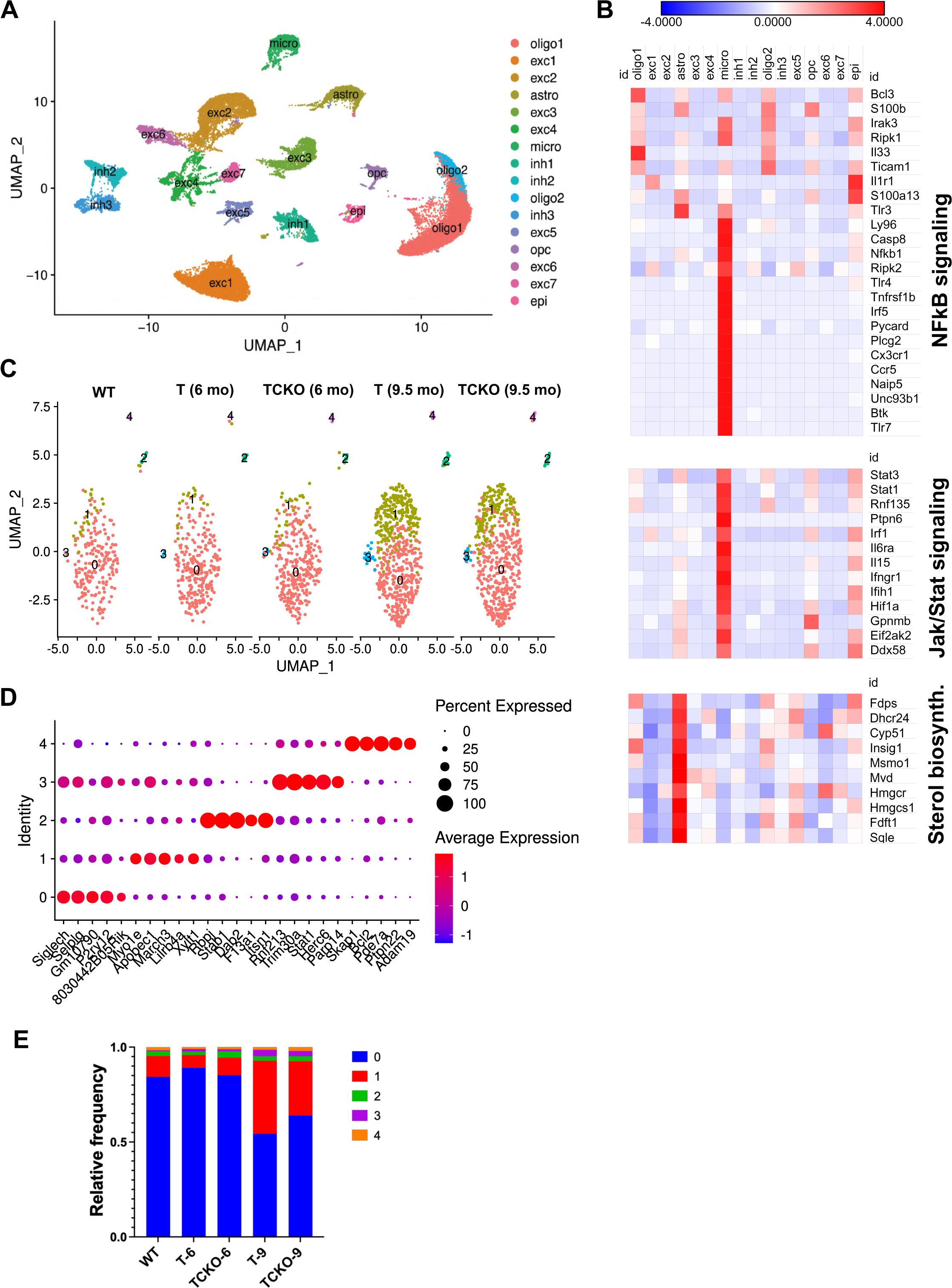
Single nuclei-RNAseq reveals microglia-specific changes resulting from Ch25h deficiency in PS19 mice. A) UMAP plot showing 16 distinguishable clusters (1–16), from snRNA-seq of hippocampal tissue from 9.5-month-old male mice (n = 5 pooled per group) with corresponding cell types identified using known cell markers (exc: excitatory neurons, inh: inhibitory neurons, astro: astrocytes, micro: microglia, oligo: oligodendrocytes, opc: oligodendrocyte precursor cells, epi: epithelial cells). B) Average expression of genes related to NF-kB sugnaling, JAK/STAT signaling and sterol biosynthesis in each cell-type signature. C) UMAP plot of the sub-clustered microglia population showing 5 distinguishable clusters (0–4) identified in WT (9.5-month old), T (6 and 9.5-month old), and TCKO mice (6 and 9.5-month old). D) Dot plot showing percentage of expression (dot size) and average gene expression levels (color intensity) of five of the most expressed specific markers in each microglia subcluster. E) Relative frequency of microglia subclusters in each experimental group.

To understand how the inflammatory genes in microglia influence neurodegeneration, the microglia cluster were subclustered into five subclusters (cluster 0-4) of microglia in WT, T and TCKO mice (Figure 8C, Figure 8D, Supplementary Table IX). Most of the nuclei from our samples were grouped in cluster 0, identified by high mRNA expression of markers (e.g., *Cxcr3* and *P2ry12*) expressed in a microglia transcriptomic state commonly referred to as “homeostatic” (Figure 8D, Supplementary Table IX). Microglia nuclei cluster 0 distribution in T and TCKO mice at 6-months old were comparable with 9.5-month old WT mice (Figure 8C, 8E). By 9.5 months of age, however, a marked increase of nuclei from cluster 1 was identified in T and TCKO mice relative to WT (Figure 8C, 8E). Cluster 1 is characterized by high expression of mRNA for markers (e.g., *Apobec, Apoe, Axl, Cd9, Lpl*) typical of the transcriptomic signature referred to as microglial neurodegenerative (MGnD) or DAM (Figure 8D, Supplementary Table IX). An age-dependent increase of nuclei grouped in cluster 3 characterized by interferon signaling genes (e.g., *Stat1, Ifit2*) (Figure 8D, Supplementary Table IX) was also observed in tau-transgenic mice compared to WT (Figure 8C, 8E).

Nuclei proportion analysis for each experimental group showed that the presence of tau pathology at 9.5 months of age promoted a decrease in the percentage of nuclei of the homeostatic microglia (cluster 0) and an increase of DAM (cluster 1) and interferon (cluster 3) genes (Figure 8E, Supplementary Table X). Ch25h deficiency not only promoted an increase of the homeostatic microglia nuclei proportion, but also decreased the DAM and the interferon subclusters (Figure 8E, Supplementary Table X). These results are in agreement with other data suggesting that the main effect of deleting Ch25h from tau transgenic mice is a reduction of microglial activation (specifically, DAM or MGnD).

Next, we examined how Ch25h-deficiency alters the key pathways identified from bulk as well as single nuclei transcriptomics studies, namely A) sterol and lipid metabolism, B) NF-kB and Jak/Stat signaling and C) leukocyte activation.

A) **Ch25h-deficiency in tau-transgenic mice reduces lipids associated with inflammation.** To explore the impact of Ch25h deficiency in cholesterol and lipid metabolism, levels of desmosterol, free cholesterol, and other lipids were quantified in the cortex (Figure S6). Although the bulk-RNA seq analysis in the hippocampus showed down-regulation of cholesterol biosynthetic enzymes in T mice respect WT mice that were restored in TCKO mice, levels of desmosterol (Figure S6A) and free cholesterol (Figure S6B) in the cortex were unaffected. From our bulk lipidomic analysis (Supplementary Table VI) several lipid classes were unchanged. However, we identified 3 subclusters (a-c) of lipids changing between WT, T and TCKO mice (Figure S6C). In cluster ‘a’, we observed a significant reduction in the levels of different cardiolipins (CL) and lyso-cardiolipins (LCL) species in T and TCKO mice relative to WT mice. Cardiolipins are a class of phospholipids that maintain stable mitochondrial membranes. Reduced CL levels have also been observed in the triple transgenic AD mouse model (3xTg-AD), implicating oxidative stress and mitochondrial dysfunction (38). These changes, however, are unaffected by Ch25h loss. In cluster ‘b’, increased levels of three lyso-phosphatidylethanolamine (LPE) species, as well as one species of phosphatidylcholine (PC), lyso-phosphatidylcholine (LPC) and a cerebroside were detected in T and TCKO mice relative to WT mice. Decreased levels of phosphatidylethanolamine and phosphatidylcholine in AD brain tissue have been observed (39, 40). The elevated levels of these lipids observed in PS19 mice with or without Ch25h suggests that this is a tau-mediated phenomenon specific to PS19 mice which may not fully replicate all the lipid changes associated with AD. Cluster ‘c’ was characterized by different species of cerebroside (CBS), sphingomyelin (SM), phosphatidylinositol (PI), ceramide (CER), sulfatide (ST), phosphatidylserine (PS) and phosphatidic acid (PA). Levels of lipids from this cluster were found significantly elevated in T mice compared with WT mice and strikingly downregulated in TCKO mice, suggesting that Ch25h deficiency may selectively alter lipid composition in the brains of tau-transgenic mice.
b) **Activated microglia show activated NF-kB and Jak/Stat signaling** Recent reports have indicated that the nuclear factor kappa-light-chain-enhancer of activated B cells (NF-kB) in activated microglia exacerbates neurodegeneration in PS19 mice (41). To extend our finding that two transcription factors of the NF-kB family seemed to be differentially expressed in TCKO and T mice, we immunostained for phospho-p65 subunit of NF-kB, an activated subunit of the NF-kB transcription factor and the microglia marker Cd11b (Figure 7B). Consistent with the transcriptomics data, we found a dramatic increase in area covered by p-p65 NF-kB immunoreactivity within Cd11b-positive cells in the hippocampus of T compared to WT mice. CD11b-specific p-p65 NF-kB immunoreactivity was significantly decreased in TCKO mice relative to the T mice (Figure 7C). No differences were observed in quantification of p-p65 NF-kB immunoreactivity in GFAP-positive astrocytes (Figure 7D). Since the pro-inflammatory cytokine IL1β is a downstream target of NF-kB, we also measured IL1ý mRNA levels and observed a significant reduction in IL1ý mRNA relative expression levels in TCKO mice in comparison with T mice (Figure 7E). We previously identified an augmented secretion of IL-1β in mouse primary microglia treated with LPS which was attenuated in Ch25h-deficient microglia (24). Signal transducer and activator of transcription 3 (Stat3) is a key regulator of the JAK/STAT signaling pathway and was markedly reduced in our bulk transcriptomic analysis. A previous report showed that Stat3 is highly phosphorylated in activated microglia and exacerbates neurodegeneration in PS19 mice (42). Consequently, we immunostained for phosphorylated Stat3 (p-Stat3) and Cd11b in the hippocampus of WT, CKO, T and TCKO mice (Figure 7F). Consistent with previous findings, p-Stat3 immunoreactivity was significantly increased in microglia of T mice and decreased in microglia of TCKO mice (Figure 7G). Immunoreactivity of p-STAT3 in astrocytes remained unchanged (Figure 7H). Taken together, our transcriptomic analyses of DEGs and identification of pathways altered by Ch25h deficiency strongly suggests that 25-HC is a critical positive modulator of neuroinflammation in PS19 mice. Therefore, we examined if the reduced inflammatory response in microglia observed in tau-transgenic mice lacking Ch25h is reproducible *in vitro* in cultured primary microglia. To this end, we investigated the inflammatory response of mouse primary microglia from WT and CKO mice treated with pre-formed fibrils (0.5μm) of human recombinant tau-441 (2N4R) P301S mutant protein (Figure 9). First, we tested the ability of Ch25h-deficient mouse primary microglia to phagocytose ATTO 488-conjugated preformed-fibrils of P301S tau after 2 hours of treatment (Figure 9A). Our results indicate that CKO and WT microglia phagocytose the tau fibrils similarly (Figure 9B), as a comparable amount of tau fibrils was found in CD68^+^ phagolysosomes in both cell types (Figure 9C). Next, we tested the effect of unlabeled pre-formed tau fibrils on the relative expression of genes related to inflammatory response in mouse primary microglia. After 24 h of treatment, no changes in Ch25h expression (Figure 9D) were identified in WT mouse primary microglia treated with tau fibrils relative to the control. While tau fibrils induced the expression of IL1β(p<0.01) in WT microglia, a negligible increase was detected in CKO microglia (Figure 9E). However, tau fibrils did not alter expression of TNFα expression in either cell type (Figure 9F). Interestingly, CKO mouse primary microglia showed upregulated levels of expression of the anti-inflammatory cytokine, IL-10, compared to WT mouse primary microglia (p<0.05, Figure 9H). Furthermore, tau fibrils elicited increased expression of Cxcl10 (p<0.01) that was markedly reduced (p<0.05) in CKO microglia treated with tau fibrils (Figure 9G). Similar suppression of Cxcl10 expression in TCKO mice relative to T mice was observed by bulk transcriptomic analysis (Supplementary Table IV). These *in vitro* data suggest that tau fibrils induce a proinflammatory response that is suppressed in the absence of Ch25h and 25HC, similar to previous reports (20, 24). These *in vitro* data suggest that tau fibrils induce a proinflammatory response that is suppressed in the absence of Ch25h and 25HC, similar to previous reports (20, 24).
C) **Ch25h deficiency reduces the T-cell infiltration observed in PS19 mice.** Infiltration of T-cells mediated by disease-associated microglia has been reported to play a crucial role in inducing neurodegeneration in PS19 mice (43). Pharmacological depletion of microglia reduced the number of T-cells in the brain and T-cell depletion resulted in decreased neurodegeneration in PS19 mice (43). Based on our bulk transcriptomic analysis of the hippocampus, genes clustered in the GO term ‘leukocyte activation’ were found downregulated in TCKO mice compared with T mice. Considering the reduced microgliosis observed in TCKO mice, we immunostained mouse brain sections for the T-cell specific receptor, CD3 (Figure 10A). Consistent with the above mentioned report (43), a significant increase in the percentage of area covered by CD3 immunoreactivity (Figure 10B) was found in the hippocampus of T mice compared to the controls (p<0.01), as well as TCKO mice (p<0.05). These results demonstrate that upregulation of Ch25h/25-HC may contribute to microglia-mediated T-cell infiltration in the PS19 mouse model.

**Figure 9:**
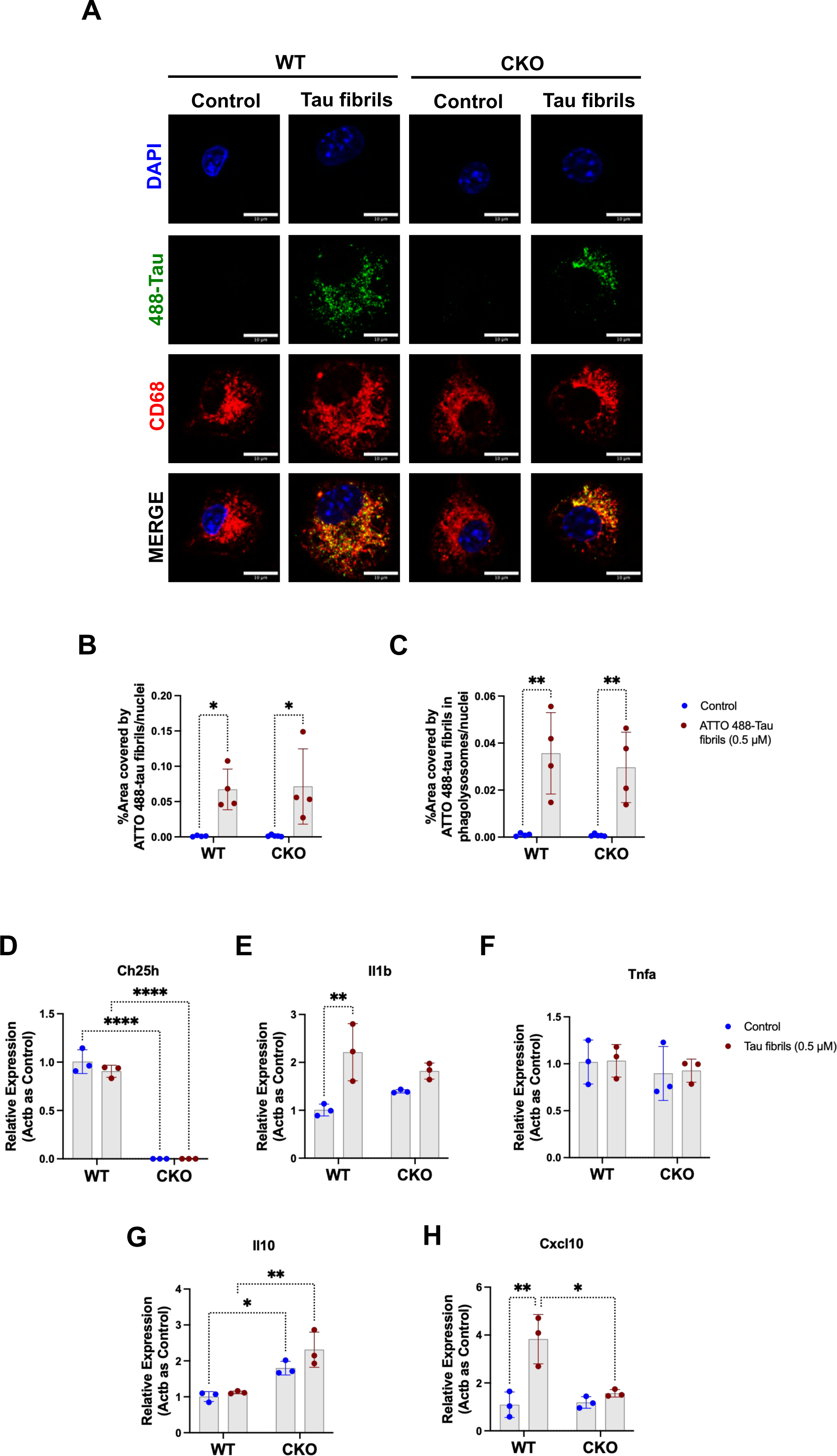
Ch25h expression controls microglial inflammatory response induced by tau fibrils. Mouse primary microglia from WT and CKO mice were treated with vehicle or 0.5μM ATTO 488-conjugated tau preformed-fibrils (green) for 2 hours. Cells were co-stained with CD68 (red) and DAPI (blue). Phagocytosis of tau fibrils was evaluated by confocal microscopy (scale bar 10 µm). (A). Quantification of the percentage area of ATTO 488-conjugated tau fibrils (green) was conducted (B) as well as the localization in CD68^+^ phagolysosomes (C). Data was normalized to number of nuclei. Expression of *Ch25h* (D), *Il1b* (E), *Tnfa* (F), *Il10* (G), and *Cxcl10* (H) was assessed by qPCR after 24 h of treatment of microglia with unlabeled tau pre-formed fibrils. Two-way ANOVA with Tukey’s post hoc test was used for the statistical analysis. *p<0.05, **p<0.01, ****p<0.0001.

**Figure 10:**
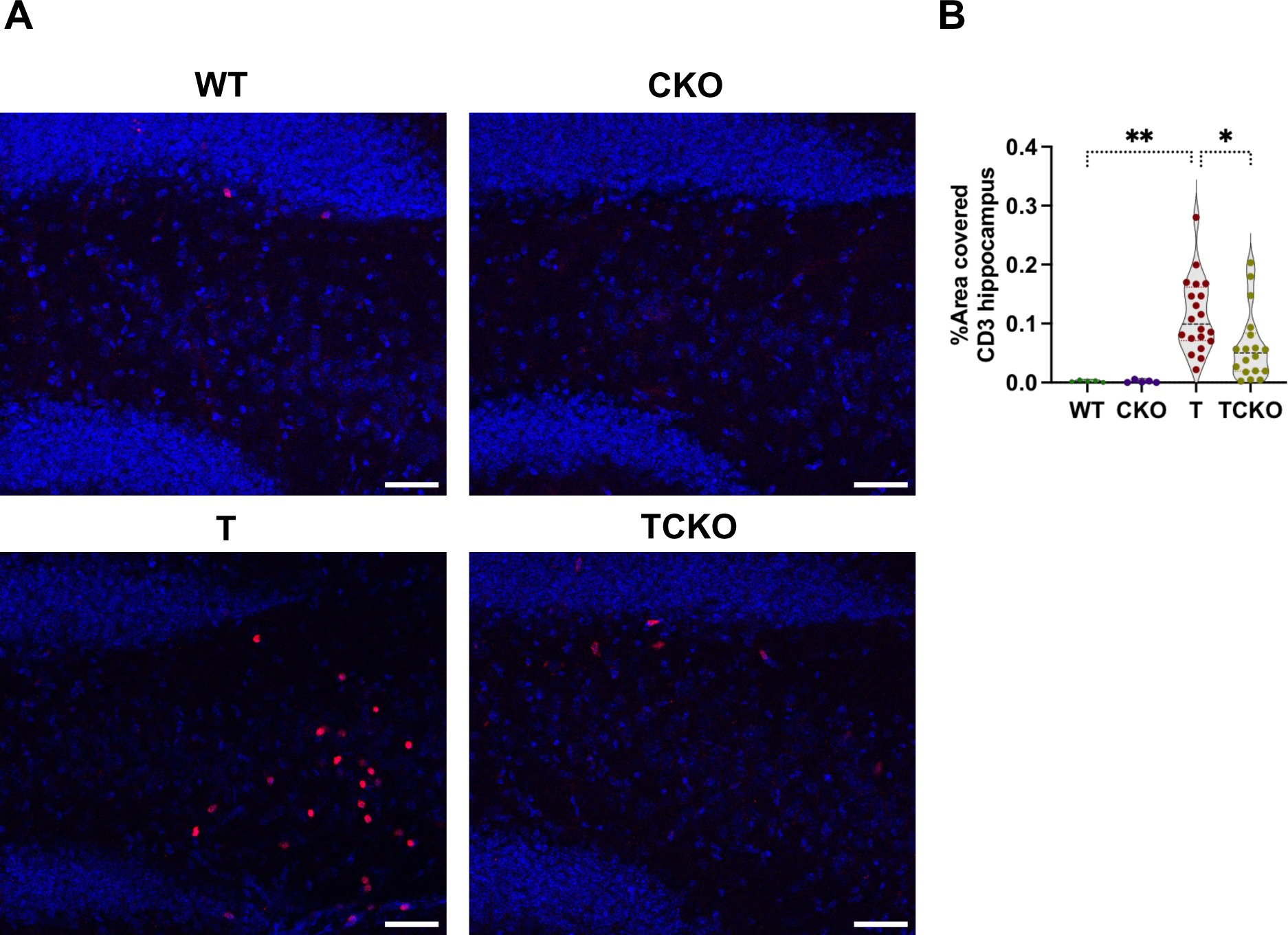
Ch25h deficiency reduces brain T-cell infiltration in PS19 mice. (A) Representative images of immunostaining of brain sections for the pan-T cell antigen, CD3 in the dentate gyrus region co-stained with DAPI in 9.5-month old wild type (WT, n=5), Ch25h KO (CKO, n=5), PS19 (T, n=20) and PS19/Ch25h KO (TCKO, n=20) (B) Quantitation of the percent area covered by CD3 in each group. One-way ANOVA with Tukey’s post hoc test (two-sided) was used for all statistical analysis. *p<0.05, **p<0.01, ***p<0.001, ****p<0.0001. Scale bar 50 µm.

## DISCUSSION

In this study, we confirm that by 9.5 months of age PS19 mice manifest significant tau accumulation, neuroinflammation (microgliosis and astrogliosis), atrophy and neuronal loss in the hippocampus and entorhinal/piriform cortex. We now report that all of these pathological hallmarks are markedly reduced in PS19 mice lacking Ch25h. We also confirm that CH25H is overexpressed by microglia in AD brain tissue as well as in the PS19 mouse model of tauopathy.

We show that increased expression of Ch25h and the concomitant increase in 25-HC levels in the brains of PS19 mice occurs after six months of age and may reflect exacerbated microglial activation. Interestingly, elevated levels of oxysterols including 25-HC were detected in late-stage AD brains (44). 25-HC is an oxysterol with potent immunomodulatory effects (45). We and others have previously shown that Ch25h is responsible for the production and secretion of 25-HC by microglia (14, 24) and that microglia from Ch25h knockout mice produce no detectable levels of 25-HC under basal conditions or when stimulated with LPS (35).

Several transcriptomic studies have also demonstrated that expression of Ch25h is limited to microglia in the CNS (22). Conversion of homeostatic microglia into DAM is a progressive change that occurs through a TREM2-independent stage (DAM1) followed by a Trem2-dependent stage (DAM2) characterized by increased expression of genes related to microglial phagocytic activity as well as lipid metabolism (e.g. *Ch25h*) (4). Our results show PS19 mice deficient in Ch25h and 25-HC have decreased expression of several DAM genes (e.g. *Apoe*, *Trem2*, and *Clec7a*) without fully restoring the homeostatic state of the cells. Thus, by simply eliminating one non-essential DAM gene, Ch25h, much of the neurodegeneration observed in aged PS19 mice was prevented similar to the effect observed by deleting *Trem2*, *Apoe* or depleting microglia in PS19 mice (7, 8, 10). Expression of Ch25h is controlled by Trem2 (12, 46) and our transcriptomic results suggested that Ch25h may amplify Trem2 signaling, suggesting that 25-HC is a lipid effector that can modulate inflammatory functions of microglia. In agreement with a previous report (47), we identified an exaggerated inflammatory gene signature characterized by GO terms such as ‘positive regulation of cytokine production’, ‘leukocyte activation’, and ‘inflammatory response’. The absence of Ch25h in PS19 mice strikingly attenuated DEGs related to these same inflammatory pathways supporting a role for Ch25h in triggering neuroinflammation through 25-HC secreted by microglia leading to age-dependent neurodegeneration. Specifically in the context of regulation of cytokine production, NF-κB signaling was among the top altered immune response pathways that potentially mediates the proinflammatory effects of 25-HC, and gene network analysis suggested a strong connection between NF-kB pathway-related genes and IL-1β production. Consistently, reduced phosphorylation of p65 NF-kB subunit and lL-1β expression was observed in Ch25h-deficient PS19 mice. In a recent study by Jiang et al. (48) phosphorylated-tau was shown to induce IL-1β expression through activation of the MyD88-NF-κB pathway and NLRP3 inflammasome in a similar tauopathy mouse model (48). Deletion of MyD88 or the inflammasome adaptor protein ASC in these mice blocked neuroinflammation, reduced p-tau levels and improved cognition. In another recent report, inactivation of NF-kB in microglia of PS19 mice partially restored microglial homeostasis, decreased tau seeding and spreading and protected against spatial learning and memory deficits (41). We have previously shown that treatment of primary mouse microglia with the TLR4 agonist LPS markedly upregulates Ch25h expression and 25-HC secretion in addition to cytokine overproduction (24). Secretion of IL-1β was augmented by 25-HC treatment of primary microglia and reduced in Ch25h-deficient microglia. 25HC-mediated augmentation of IL-1β secretion was due to enhanced caspase 1 activity via NLRP3 inflammasome activation (24). We corroborated that wildtype, but not Ch25h-deficient, mouse primary microglia stimulated by P301S tau fibrils express higher levels of IL-1β mRNA. Moreover, Ch25h-deficient microglia express increased levels of the anti-inflammatory protein, IL-10, but decreased levels of the chemokine, Cxcl10. Cxcl10 is a key interferon gamma (INF-γ) -inducible chemokine that has been previously reported to be stimulated by tau fibrils in mouse primary microglia (49), as well as in PS19 mice where it may facilitate T-cell infiltration into the brain (43).

T-lymphocyte infiltration into the brain has been previously reported in AD patients and has been associated with impaired cognition (50, 51). Recent reports in two different mouse models of tauopathy (THY-tau22 and PS19 mice) suggest that exhausted activation of microglia is crucial to promoting the T-cell infiltration into brain which exacerbates neuroinflammation and neurodegeneration and impairs cognition (43, 52). 25-HC is metabolized by Cyp7b1 to 7α,25-dihydroxycholesterol, which is a high affinity chemoattractant for leukocytes expressing the G-protein coupled receptor, Gpr183 (also called EBI2) (53). Our finding of reduced T-cells in the hippocampus of Ch25h-deficient PS19 mice may be due to decreased chemokine secretion from activated microglia (e.g. Cxcl10) and a lack of the well characterized production of 7α,25-dihydroxycholesterol. In a mouse model of experimental autoimmune encephalomyelitis (EAE), expression of Gpr183 was demonstrated in a variety of T cells and was shown to be required for the encephalitogenic T-cell infiltration into the brain (54). We speculate that in PS19 mice increased expression of Ch25h results in increased 25-HC and 7α,25-dihydroxycholesterol production leading to increased CNS infiltration of leukocytes, further contributing to inflammation-induced neurodegeneration.

The fact that Ch25h-deficient PS19 mice also show less astrogliosis, suggests that 25-HC secreted by microglia may contribute to the abnormal astrocyte function observed in neurodegenerative disorders like AD. Among the genes expressed in astrocytes, loss of Ch25h restored the expression of sterol biosynthesis genes which were significantly downregulated by tau pathology in PS19 mice. Downregulation of genes necessary for cholesterol biosynthesis in 7 and 9-month old PS19 mice has been recently reported (55). Abnormal cholesterol metabolism has also been implicated in AD, however both increased and decreased levels of cholesterol and its metabolites have been reported (56–58). Decreased expression of the cholesterol biosynthetic enzymes and the cholesterol biosynthetic intermediate, lanosterol, without significant changes in free cholesterol levels has also been recently reported in AD brain tissue (59). In our study, we did not detect changes in free cholesterol or cholesterol precursors in the cortex of PS19 mice. Our snRNAseq analysis shows however that the cholesterol biosynthetic enzymes are expressed mainly in astrocytes which are critical for cholesterol biosynthesis in the brain. The underlying mechanism by which 25-HC suppresses sterol biosynthesis genes likely involves its ability to inhibit the proteolytic processing of sterol response element-binding protein 2 (SREBP2), a master transcription factor that controls the transcription of genes encoding cholesterol biosynthetic enzymes (60). We have recently shown that 25-HC inhibits cholesterol synthesis and promotes cholesterol esterification in primary mouse astrocytes (35). Interestingly, astrocyte-specific deletion of SREBP2 has been reported to markedly impair brain development, neuronal function, and behavior in mice (61). A recent study using an in vitro model of tau entry into neurons showed that depletion of cholesterol from the plasma membrane or addition of 24(S)-HC, promoted tau entry into the cytosol whereas 25-HC reduced it (33). However, we did not detect any effect of Ch25h-deficiency on tau seeding or spreading in vivo. Thus, we speculate that 25-HC inhibits SREBP2-mediated gene expression leading to a suppression of cholesterol biosynthesis in astrocytes, which in turn may contribute to depletion of cholesterol in neurons, resulting in impaired neuronal function and possibly neurodegeneration in PS19 tauopathy mice.

Abnormal levels of various phospholipids and sphingolipids have also reported in AD (39, 40, 58, 62). Elevated levels of ceramides and galactosyl-ceramides have been reported in the middle frontal gyrus of AD patients (63). In our study, increased levels of sphingomyelins, cerebrosides (glucosyl- and galactosyl-ceramides) and ceramide were observed in PS19 mice suggesting a dysregulation of sphingolipid metabolism mediated by tau accumulation which was restored by the loss of Ch25h. Accumulation of ceramides in the brain can be stimulated by cytokines and oxidative stress, and ceramides can amplify neuroinflammation and promote apoptosis (64). It is tempting to speculate that the reduction in sphingolipids observed in Ch25h-deficient PS19 mice is related to the reduction in neuroinflammation and neurodegeneration.

Collectively, our data show that 25-HC produced by activated microglia contributes to tau-mediated neuroinflammation and neurodegeneration in PS19 mice. It will be important in future studies to assess the effects of 25-HC on behavior in PS19 mice. It is likely that 25HC exerts synergistic effects with other cytokines and (or) chemokines as well as lipid mediators, resulting in age-dependent proinflammatory actions as well as recruitment of other immune cells into the brain. 25-HC may also elicit autocrine and paracrine effects to inhibit sterol biosynthesis necessary for normal neuronal function and viability. Thus, inhibitors of either Ch25h expression or enzyme activity may offer a novel strategy for reducing the neuroinflammation and neurodegeneration which characterizes AD and other tauopathies.

## MATERIAL AND METHODS

### Animals

The PS19 transgenic mouse model overexpresses 1N4R human microtubule-associated protein tau (*MAPT*) driven by the PrP promoter, containing the P301S mutation that causes a familial form of frontotemporal dementia (25). PS19 tau transgenic mice (#008169; The Jackson Laboratory) were backcrossed and maintained on a C57BL/6 background. Ch25h knockout mice (B6.129S6-*Ch25h^tm1Rus^*/J) were purchased from The Jackson Laboratory (#016263). This strain was backcrossed to C57BL/6J for more than 10 generations by the donating laboratory. The PS19 (T) mice were crossed to the Ch25h knockout (CKO) mice to generate wild type (WT), Ch25h KO (CKO), T and TCKO mice. Only female mice were used for analysis in this study. All animal procedures and protocols were approved by the Institutional Animal Care and Use Committee at the Washington University School of Medicine.

### Tissue collection

At 9.5 months of age, mice were anesthetized with pentobarbital (i.p. 100 mg/kg) and sacrificed by cardiac perfusion with cold Dulbecco’s phosphate buffered saline (DPBS). The left hemisphere was collected and fixed in 4% paraformaldehyde overnight before being transferred to 30% sucrose and stored at 4°C until they were sectioned coronally at 40-μm on a freezing sliding microtome (Thermo Scientific, HM430). Sections were stored in cryoprotectant solution at –20°C until use. Hippocampus and cortex were dissected from the right hemisphere and kept at -80°C until used for RNA isolation and biochemical analysis.

### Volumetric Analysis

Every sixth brain section (240 mm between sections) starting rostrally at –1.3 mm to the dorsal end of the hippocampus at bregma –2.8 mm was mounted on slides and allowed to dry overnight at room temperature. The following day, mounted sections were stained with 0.1% Sudan black in 70% ethanol at room temperature for 20 minutes, then washed in 70% ethanol 3 times for 1 minute. The sections were rinsed in Milli-Q water and coverslip was placed with Fluoromount-G (Southern Biotech, 0100-01). Slides were imaged with the NanoZoomer 2.0-HT system (Hamamatsu Photonics). Hippocampus, posterior lateral ventricle and entorhinal/piriform cortex areas were blinded to genotype and measured using the NDPview.2 software (Hamamatsu Photonics). The volume for each region of interest was calculated using the following formula: volume= (sum of areas) x 0.24 mm.

### Neuronal layer thickness

Three sections (bregma -1.34, -2.3, -3.26 mm) per mouse were stained with DAPI (Invitrogen; D1306) and mounted with Fluoroshield. Images were acquired using the Zeiss Axio Scan 7 (Zeiss). The thickness of the CA1 pyramidal cell layer, dentate gyrus granular cell layer and pyramidal layer from the piriform cortex were blinded to genotype and measured by drawing a scale perpendicular to the cell layer at two spots in all three slices and taking the average value for each mouse, using the Zen lite software (Zeiss).

### Immunohistochemistry

Three sections (bregma -1.38, -2.34, -3.3 mm) per mouse were washed 3 times in TBS for 5 minutes and blocked in 0.3% hydrogen peroxide for 10 minutes. After washing, sections were blocked in 3% milk in TBS with 0.25% Triton X-100 (TBSX) for 30 minutes. AT8 primary antibody (mouse monoclonal-biotin, 1:500; Thermo Fisher Scientific, MN1020B) was diluted in 1% milk in TBSX, and the sections were incubated at 4°C overnight. The next day, sections were washed in TBS and incubated at room temperature in VECTASTAIN Elite ABC system (Vector Laboratories, PK-6100) for 1 hour, followed by another washing step. Sections were developed in DAB solution (Sigma, D5905) for 10 min, washed, and mounted on slides. After drying overnight, the slides were dehydrated in increasing ethanol concentrations (from 50%-100%) followed by xylene and coverslipped with Cytoseal 60 (Thermo Fisher Scientific, 8310). Slides were scanned on the NanoZoomer 2.0-HT system. Images were processed and AT8 percentage of area covered in the hippocampus and piriform/entorhinal cortex were quantified in the Fiji software version 2.3.0, using macro in batch mode.

### Immunofluorescence

Free-floating sections (3 per mouse, 960 μm apart from each other) were washed 3 times, 5 min each into 12-well plates with Netwell inserts (Corning) containing TBS. Tissues were blocked and permeabilized in 5% normal goat serum (NGS) and 3% bovine albumin serum (BSA) in TBSX for 1 hour at room temperature (5% donkey serum in 0.5% TBSX for Trem2 immunofluorescence). Tissue was then incubated overnight at 4°C with primary antibodies CD68 (rat monoclonal, 1:400; Biorad, MCA1957), Iba1 (rabbit polyclonal, 1:1000; FUJIFILM Wako’s, 019-19741), GFAP (chicken polyclonal, 1:1000; Abcam, ab4674), Clec7a or Dectin-1 (rat monoclonal, 1:100; Invivogen, mabg-mdect), Trem2 (sheep polyclonal, 1:400; R&D systems, AF1729), ApoE (mouse monoclonal HJ6.3, 1:1000; from D. Holtzman); Tmem119 (rabbit polyclonal, 1:250; Cell Signaling, 90840S), P2ry12 (rat monoclonal, 1;100; Biolegend, 848002), Cd11b (rat monoclonal, 1:400; BioLegend, 101202), p-STAT3 (rabbit polyclonal, 1:500; Cell Signaling, 9145S), p-p65 NF-kB (rabbit polyclonal, 1:3000; Cell Signaling, 3033S), MC1 antibody (mouse monoclonal, 1:500: gift from Peter Davis), CD3 (rat monoclonal, 1:500; Invitrogen, 14-0032-82). The following day, sections were washed 3 times in TBS, followed by 1 hour incubation at room temperature in secondary antibodies diluted in TBS (Invitrogen, 1;1000): goat anti-rabbit Alexa 488 (A32731), goat anti-rat Alexa 555 (A21434), goat anti-mouse Alexa 647 (A32728), goat anti-mouse Alexa 555 (A32727), goat anti chicken Alexa 647 (A21449), donkey anti-sheep Alexa 555 (A21436), donkey anti-rat Alexa 488 (A21208).

Sections were washed with TBS 3 times and mounted on Fisherbrand™ Superfrost™ Plus Microscope Slides (Fisher Scientific,12-550-15) and then coverslipped using Fluoroshield mounting medium. For quantification purposes, hippocampus and entorhinal/piriform cortex were scanned in the Nikon Spinning Disk and analyzed with Fiji software version 2.3.0, using macro in batch mode. After analysis, representative images were acquired on an LSM 880 II (Zeiss) microscope. For P2ry12 quantification, a surface analysis was conducted using Imaris 9.5 software (Bitplane).

### Glia morphology analysis

Three z-stacks (10 μm, 960 μm a er) of GFAP and Iba1 immunostained sections were acquired on an LSM 880 II (Zeiss) with a 63X objective. Glia morphology was analyzed using the filament tracer in Imaris 9.5 software (Bitplane) as reported previously (65). Number of branches points and process length per cell were plotted. For microglia analysis the following settings were used: Number of process length largest diameter 9.00 μm, seed points 2.200 μm; remove seed points around starting points; and diameter of sphere regions: 15 μm. The following settings were applied for astrocyte morphology analysis: Number of process length largest diameter 8.00 μm, seed points 2.000 μm; remove seed points around starting points; and diameter of sphere regions: 8 μm

### Synaptic loss analysis

Immunofluorescence for synaptic markers Synapsin 1/2 (guinea pig polyclonal, 1:500; Synaptic Systems, 106004) and PSD95 (rabbit polyclonal, 1:200; Thermo Fisher Scientific, 51-6900) were performed as previously described (66), using the secondary antibodies goat anti-guinea pig Alexa 488 (1:1000; Invitrogen, A11073) and goat anti-rabbit Alexa 594 (1:1000; Invitrogen, A11012). Images were acquired at 63x in the stratum lucidum of CA3 region on an LSM 880 microscope with AiryScan detector (Zeiss). A spot analysis was conducted in Imaris 9.5 software. Spots were detected using an automated background subtraction and spot settings detection for each channel (Synapsin spots: *x*-*y* size of 0.9 μm; PSD95 spots: *x*-*y* size of 0.5 μm).

### Immunofluorescence of human tissue

Human brain sections were obtained from the Banner Sun Health Institute. Demographic information is detailed in Supplementary Table I. For human tissue, 50-micron free-floating sections were washed in TBS followed by antigen retrieval performed with 10 mM sodium citrate buffer, pH 6 and 0.05% Tween at 60 °C for 30 min. Sections were blocked with 5% goat serum in TBSX (0.5% Triton X-100). Antibodies were incubated in blocking solution at 37°C for 2h: Iba (rabbit polyclonal), CH25H (mouse monoclonal, 1:150; from the hybridoma supernatant kindly provided by Dr. David Russell from UT South-Western), GFAP (chicken polyclonal) and anti-mouse biotin AT8 (mouse monoclonal). For sections stained with AT8 we use Streptavidin Alexa 488 (1:500, Thermo Fisher Scientific, S32354). The sections stained with IBA1, GFAP and CH25H, we used the secondary antibodies (Invitrogen, 1;1000): goat anti-chicken Alexa 488 (A11039), goat anti-rabbit Alexa 555 (A32732), goat anti-mouse Alexa 647 (A32728). Three images (z-stacks 16 µm) per tissue were acquired on an LSM 880 microscope with at 20x magnification. 3D-surfaces from CH25H, IBA1 and GFAP immunoreactivity were generated using a semi-automated pipeline based on MATLAB in Imaris 9.5 software (Bitplane). Total volume of CH25H was plotted. Next, surfaces of CH25H colocalizing with either IBA1 or GFAP surfaces were created, and the colocalization volume of CH25H and IBA1, and colocalization volume of CH25H and GFAP were quantified.

### RNAscope

RNAscope® in situ hybridization assay was performed according to manufacturer’s instructions for fixed-frozen free-floating sections (Advance Cell Diagnostics (ACD), Hayward, CA, USA). Incubation was performed on the HybEz™ hybridization system (ACD). Ch25h Mouse RNAscope® probe was purchased from ACD (Cat No. 424561), as well as the negative (Cat. No. 310043, ACD) and positive (Cat. No. 313911, ACD) control probes. Detection was performed using RNAscope® 2.5 HD Assay-RED. Following *in situ* hybridization, the sections were processed for immunofluorescence as briefly described using the rabbit anti-Iba-1 antibody with corresponding Goat anti-rabbit Alexa Fluor^®^ 488 secondary antibody. DAPI solution was added with the secondary antibody for counterstained nuclei. Images were acquired on an LSM 880 II (Zeiss) microscope and were processed similarly in Imaris 9.5 software (Bitplane).

### RNA Sequencing and Data Analysis

Total RNA was isolated from mouse hippocampus with the *Quick*-RNA Miniprep Plus Kit (Zymo Research, R1058) and RNA integrity was determined using Agilent Bioanalyzer. Library was prepared with 10ng of starting RNA with a Bioanalyzer RIN score greater than 8.0. ds-cDNA was generated using the SMARTer Ultra Low RNA kit for Illumina Sequencing (Takara-Clontech) per manufacturer’s protocol. cDNA was fragmented using a Covaris E220 sonicator using peak incident power 18, duty factor 20%, cycles per burst 50 for 120 seconds. cDNA was blunt ended, had an A base added to the 3’ ends, and then had Illumina sequencing adapters ligated to the ends. Ligated fragments were then amplified for 12-15 cycles using primers incorporating unique dual index tags. Fragments were sequenced on an Illumina NovaSeq-6000 using paired end reads extending 150 bases. Basecalls and demultiplexing were performed with Illumina’s bcl2fastq software and a custom python demultiplexing program with a maximum of one mismatch in the indexing read. RNA-seq reads were then aligned to the mouse genome (Ensembl release 76 primary assembly) along with additional human *MAPT* gene with STAR version 2.5.1a1 (67). Gene counts were derived from the number of uniquely aligned unambiguous reads by Subread:featureCount version 1.4.6-p52 (68).

All gene counts were then imported into the R package EdgeR (69) where TMM normalization size factors were calculated to adjust for samples for differences in library size. Ribosomal genes and genes not expressed in three samples with count-per-million (CPM) greater than 1 were excluded from further analysis. The TMM size factors and the matrix of counts were then fed into the R package Limma (70). Weighted likelihoods based on the observed mean-variance relationship of every gene and sample were then calculated for all samples with the voomWithQualityWeights (71). The performance of all genes was assessed with plots of the residual standard deviation of every gene to their average log-count with a robustly fitted trend line of the residuals. Differential expression analysis was then performed to analyze for differences between experimental groups and the differentially expressed genes (DEGs) were defined as those with fold-change greater than 1.5 at p-value less than 0.05. The Benjamini-Hochberg false-discovery rate adjusted p-values were also determined for further analysis.

### Pathway Enrichment Analysis

Metascape (37), a web-based gene annotation and biological enrichment tool that incorporates all major biological annotation databases including biological process in gene ontology terms, was utilized for pathway enrichment analysis. One of its key features was collapsing redundant biological terms with similar set of DEGs into representative terms to help capture most significantly enriched pathways. It also analyzed the relevant transcriptional factors of those DEGs using textmining based TRRUST database (72). To help illustrate protein-protein interactions within specific enriched biological processes, the STRING data bases (73) were used to draw interactive networks of those key biological pathways using CytoScape (74).

### Gene expression by quantitative PCR (qPCR)

Expression of some genes were corroborated by synthesizing cDNA with the High-Capacity RNA-to-cDNA kit (Applied Biosystems). For the qPCR reaction mix, PrimeTime probe-based qPCR assays and PrimeTime Gene Expression Master mix (Cat. No.1055771) were obtained from IDT (Integrated DNA Technologies, Inc). qPCR reactions were run using the Fast mode on a QuantStudio™ 3 Real-Time PCR Instrument (Applied Biosystems by ThermoFisher, A28131). Data were normalized against actin (Actb) and the 2^−ΔΔCt^ method was used to calculate relative gene expression value (Relative Quantity, RQ). Relative expression of Ch25h (IDT, Assay Reference Mm.PT.58.42792394.g), Clec7a (IDT Mm.PT.58.42049707) Il1b (IDT, Assay Reference Mm.PT.58.42940223), Tnfa (IDT, Assay Reference Mm.PT.58.29509614), Cxcl10 (IDT, Assay Reference Mm.PT.58.43575827), Il10 (IDT, Assay Reference Mm.PT.58.23604055) and Actin (IDT, Assay Reference Mm.PT.58.33540333) was assessed by qPCR.

### Stereotactic intracerebral injections of AD-tau human aggregates in tau transgenic mice

AD-tau was isolated from a human AD brain at a final concentration of 5 μg/μl tau as previously described (34, 75). AD-tau preparation was diluted to 0.4μg/μl final concentration and sonicated for 30 seconds. Two-month-old T and TCKO mice were anesthetized with isoflurane and placed in a stereotaxic instrument. Mice were injected in the left hemisphere with 2.5 μl of diluted tau (0.4 μg/μl) aggregates per injection site (1 µg of tau, infusion rate 0.2 µg/µl) in the dentate gyrus (bregma: -2.5 mm; lateral: -2.0 mm; depth: -2.0 mm) and cortex (bregma: -2.5 mm; lateral: -2.0 mm; depth: -1.0 mm). Mice were monitored for 7 days after surgery and were sacrificed 45 days after the surgery.

### Quantification of oxysterols

Cortex from 6- and 9.5-month old mice were homogenized in water (9.9 mg tissue/ml) using Omni Bead Ruptor 24 (Omni International, Inc.). Oxysterols were extracted (liquid-liquid extraction) from 250 μl of homogenate after addition of 4 ng of the internal standards 24-HC-d7, 25-HC-d6, 27-HC-d5. Samples were derivatized with nicotinic acid to increase the mass spectrometric sensitivities of the oxysterols and internal standards. Quality control (QC) samples were prepared by pooling a portion of study samples and injected every 5 samples to monitor instrument performance. The sample analysis was performed with a Shimadzu 20AD HPLC system coupled to a 4000QTRAP mass spectrometer operated in positive multiple reaction monitoring mode. Data processing was conducted with Analyst 1.6.3. The quantification data were reported as pg/mg of tissue.

### Lipidomics

Lipidomic analysis was carried out at the Functional Lipidomics Core at Barshop Institute for Longevity and Aging Studies of the University of Texas Health Science Center, San Antonio, TX. Briefly, lipid extracts of cortex from 4-5 animals per group, were prepared and processed as previously described for the quantification of free fatty acids, free cholesterol, and desmosterol (35). Data processing based on the principles of shotgun lipidomics such as selective ionization, low concentration of lipid solution, and correction for differential isotopologue patterns and kinetics of fragmentation was conducted as previously described(76).

### Nuclei isolation from frozen hippocampus and Single-nuclei RNA sequencing

Frozen hippocampus from five mice per group were pooled in one sample. Nuclei isolation was conducted as described previously (77). The 10X Genomics protocol were followed, subjecting the nuclei to a droplet-based 3′ end massively parallel single-nuclei RNA sequencing using Chromium Single Cell 3′ Reagent Kits (10x Genomics). The libraries were sequenced using an Illumina HiSeq3000 (Illumina). Sample demultiplexing, barcode processing, and single-cell 3′ counting was performed using the Cell Ranger Single-Cell Software Suite (10x Genomics). The Seurat v3, MAST, and SoupX R packages were used for subsequent analysis (78–80). Contamination of cell-free RNA for each sample group was removed using SoupX. Nuclei with mitochondria content > 5% or total UMI < 300 or > 6000 were then removed in Seurat. For each group, the % of mitochondria was regressed out, gene counts normalized, and variable features identified using the SCTransform function in Seurat. The top 3000 variable genes were used to integrated experimental groups using the PrepSCTIntegration, FindIntegrationAnchors, and IntegrateData commands in Seurat. Principal component analysis (PCA) was performed on the integrated dataset and the first 30 PCs were selected for downstream analysis using FindNeighbors and RunUMAP. Clusters were identified using the FindClusters function with a granularity ranging from 0.1 to 1.2. Differential gene expression was performed using LogNormalized RNA counts using MAST. Clusters containing high mitochondrial-genome content or marker genes for more than one coarse cell type (i.e. microglia and excitatory neurons) were removed and data were reclustered using the first 15 PCs and a resolution of 0.2. For microglia subclustering, the microglial cluster was split by experimental group and re-normalized and integrated as described above. The first 15 principal components were used for the FindNeighbors and RunUMAP functions, clustering resolution was set at 0.3. Differential gene expression was obtained from LogNormalized RNA counts using MAST. Clusters containing high levels of marker gene expression for non-immune cell types were removed and data reclustered.

### Mouse primary microglia treated with tau fibrils

Mixed glia cultures were obtained from cortices of neonatal wild-type (WT) and Ch25h knock out (CKO) mice (2–3 days old) as previously described and maintained in cell culture media (DMEM + 10% heat-inactivated FBS + 1X Glutamax + 1X Sodium pyruvate and 1X Penicillin/Streptomycin, all from GIBCO) for 7 days (35). When large numbers of microglia were observed floating, the flasks were shaken at 200 rpm for 1 hour at 37°C. The floating cells were collected by centrifugation, resuspended in cell culture media. Microglia were seeded at a density of 200,000 cells/ml in 12-well plate for gene expression analysis and in 8-well chambers for assessing phagocytosis of tau fibrils. The next day, media was replaced with serum-free media (DMEM/F12 + 1X Glutamax + 1X Sodium pyruvate + 0.1% of fatty acid-free BSA and 1X penicillin/streptomycin + 1X insulin, transferrin, and selenite supplement (R&D Systems AR013) + 25 ng/ml M-CSF (GoldBio 1320-09-10)).

For tau uptake assay, WT and CKO mouse primary microglia were treated either with ATTO 488-conjugated human recombinant tau-441 (2N4R) P301S mutant protein (0.5μm) pre-formed fibrils (StressMarq Biosciences, Cat. No. SPR-329-A488) or PBS in serum-free media. After 2 hours,cells were rinsed in sterile PBS and fixed in 4% PFA for 12 minutes. Microglia were rinsed 3 times in PBS and blocked-permeabilized in 5% goat serum in TBSX. The primary antibody, anti-rat CD68 (rat monoclonal, 1:400; Biorad, MCA1957), was incubated for 2 h at 37°C and removed by rinsing the cells 3 times with TBS. Cells were incubated with goat anti-rat Alexa 555 and DAPI. Images were acquired with the LSM 880 confocal microscope at 40x (with 2x zoom). Immunoreactivity and number of nuclei were quantified in Fiji.

For assessing gene expression, microglia were treated either with endotoxin-free pre-formed fibrils of human recombinant tau-441 (2N4R) P301S mutant protein (0.5μm) expressed with a Baculovirus/sf9 system (StressMarq Biosciences, Cat. No. SPR-471) or PBS in serum-free media. After 24 hours, RNA was isolated from cells using the *Quick*-RNA Miniprep Plus Kit (Zymo Research) and cDNA synthetized as mentioned before.

### Statistics

Data are presented as mean ± SD. GraphPad Prism 9.2 was used to perform statistical analyses. Gaussian distribution was evaluated using the D’Agostino & Pearson normality test. Differences between groups were evaluated by one-way ANOVA tests with post-hoc Tukey (parametric) or Kruskal-Wallis (non-parametric) multiple comparisons tests.

### Study approval

All animal procedures and experiments were performed under guidelines approved by the animal studies committee at Washington University School of Medicine.

## AUTHOR CONTRIBUTIONS

DTR, JML, DMH, AGC, SMP: Conceived and designed research studies; DTR, JML, MRS, XH, AGC: conducted experiments, acquired data; DTR, JML, JY, JDU, XH, AGC: analyzed data; DTR, AGC, SMP: wrote the manuscript. All authors discussed the results and commented on the manuscript.

## ACKNOWLEDGEMENTS

This research was supported by funding from a grant from the National Institute on Aging of the National Institutes of Health under Award Number U19AG069701 (Project 2: AGC, JDU, SMP, and DMH) and R01AG081419 (AGC). JML was supported by career development awards from the National Institute on Aging (K08AG068611) and Alzheimer’s Association (AACSF-18-564776). We also acknowledge technical, and equipment support from the Hope Center Alafi Neuroimaging Laboratory, the Washington University Center for Cellular Imaging (WUCCI) and the Genome Technology Access Center at Washington University School of Medicine. We thank Drs. Eric Reiman, Thomas Beach, and Geidy Serrano for the human brain tissue (NIH grant 5P30AG019610). We thank the technical support for the animal breeding to Kaylee Stillwell, Ramya Changalavala, Sheryl Eveland, Ainsley Tran, Joseph Reznikov, and Rachel Schave. The authors also wish to acknowledge technical advice from members of the Holtzman, Paul and Cashikar Labs.

## SUPPLEMENTARY LEGENDS

**Figure S1:**
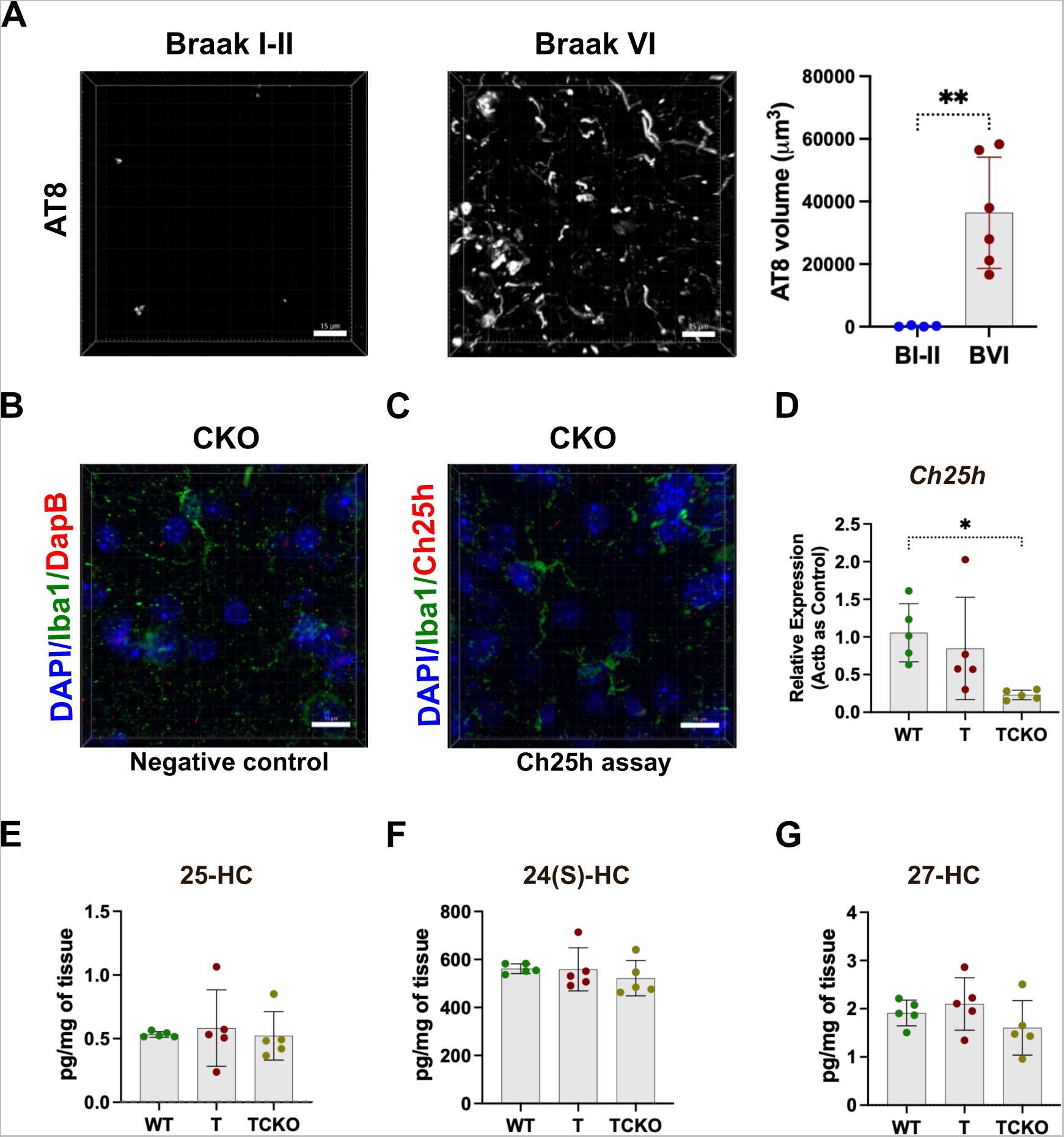
Ch25h is expressed mainly in microglia in AD and PS19 mice brains. A) Representative images of a Braak VI AD frontal cortex section stained with AT8 (phosphorylated tau) and quantification of the volumes of AT8 immunoreactivity. (B) Representative 3D Imaris images from the hippocampus of 9.5-month old female Ch25h knockout (CKO) mouse stained with DapB (B, negative control) or Ch25h assay (C) RNAscope assay, DAPI and Iba1. (D) Relative expression of Ch25h in the hippocampus of 6-months-old WT, T and TCKO female mice (n=5 for each group). Levels of 25-hydroxycholesterol (E), 24-hydroxycholesterol (F) and 27-hydroxycholesterol (G) were quantified in the hippocampus of 6-month female WT (n=5), T (n=5) and TCKO (n=5) mice. Scale bar 15 µm. Data expressed as mean ± SD. One-way ANOVA with Tukey’s post hoc test (two-sided) was used for all statistical analysis *p<0.05, **p<0.01.

**Figure S2.**
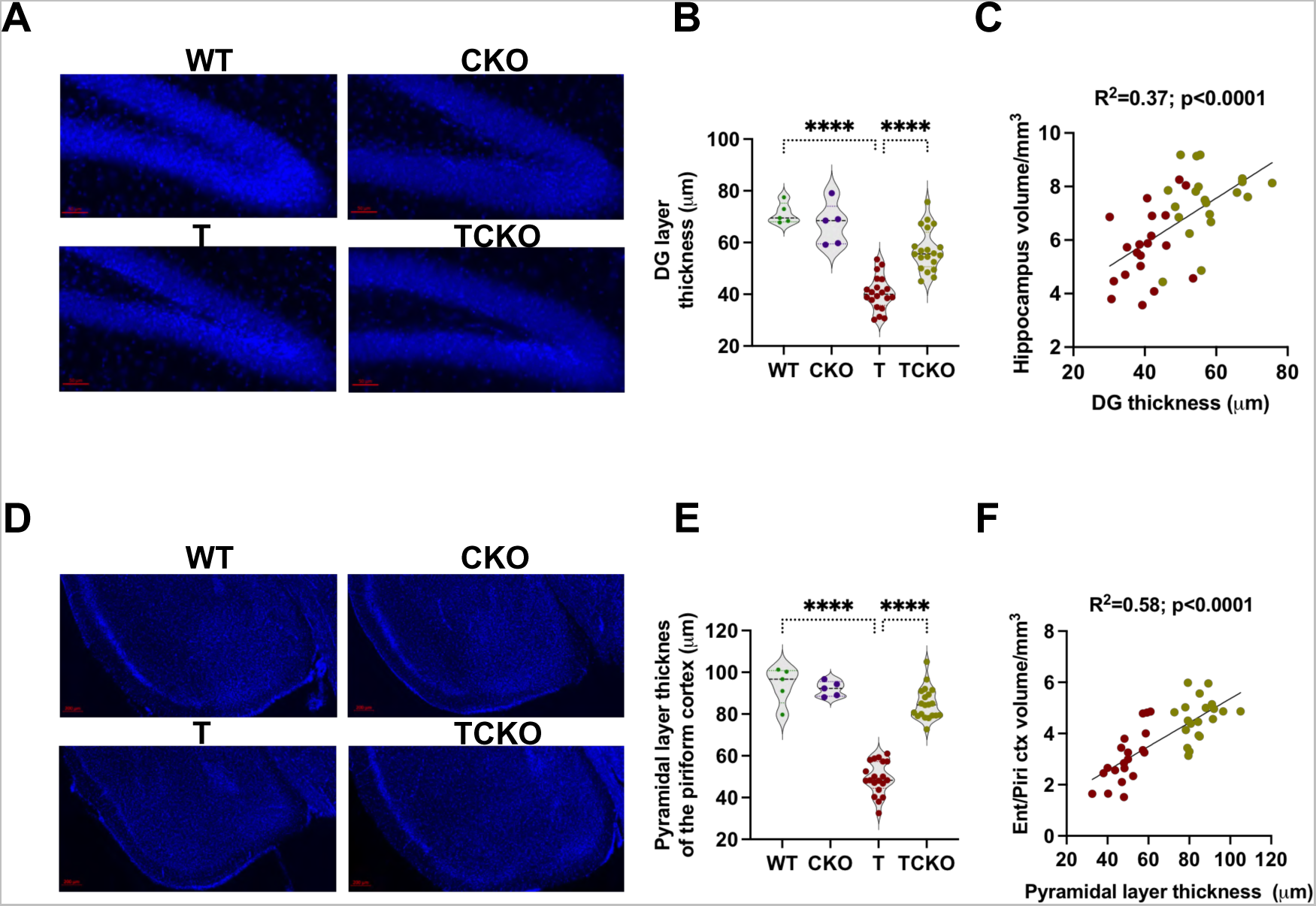
Ch25h deficiency reduces neuronal loss in PS19 mice. Representative images (A, D) and corresponding layer thicknesses (B, E) of 9.5-month old wild type (WT), Ch25h KO (CKO), PS19 (T) and PS19/Ch25h KO (TCKO) mouse brain sections stained with DAPI (Scale bar 50 µm). Thickness of the granule cell layer in DG (A, B) and the pyramidal layer in piriform cortex (D, E) were shown. Pearson correlation analysis (two sided) between DG (C) cell layer thickness with the hippocampal volume as well as the correlation between pyramidal layer of the piriform cortex (F) with the volume of entorhinal cortex are shown for tau mice - T (red) and TCKO (asparagus). Data expressed as mean ± SD, One-way ANOVA with Tukey’s post hoc test (two-sided) was used for all statistical analysis ****p<0.0001. Ent: Entorhinal cortex, Piri: Piriform cortex, DG: Dentate gyrus.

**Figure S3.**
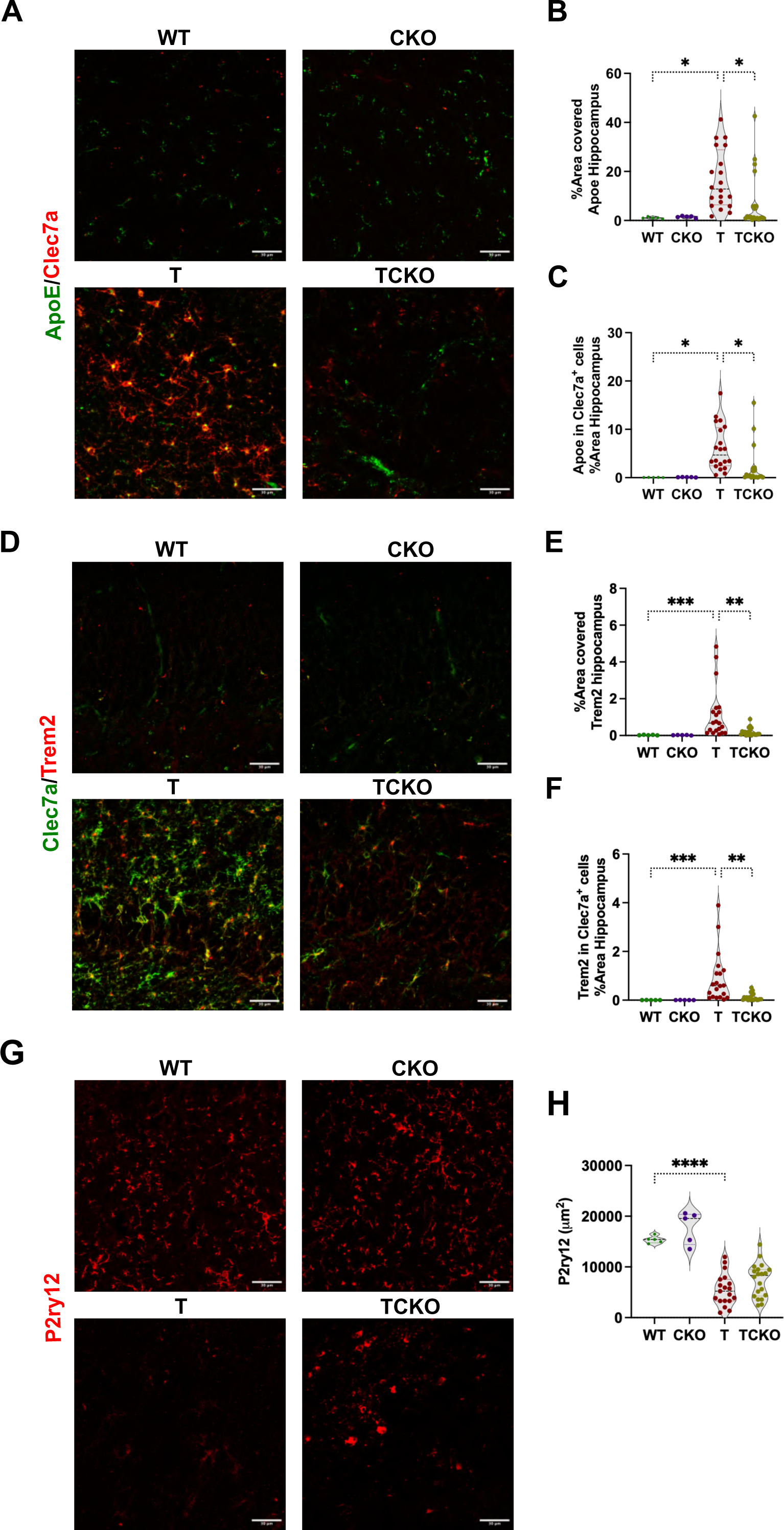
Ch25h deficiency in PS19 mice decreases DAM markers without altering homeostatic microglia. A) Representative images from a double stain of the DAM markers ApoE (green) and Clec7a (red) in 9.5-month old wild type (WT; n=5), Ch25h KO (CKO; n=5), PS19 (T; n=20) and PS19/Ch25h KO (TCKO; n=20) mouse brain sections. Percentage of area covered by ApoE immunoreactivity (B) and ApoE immunoreactivity in Clec7a positive cells (C) was quantified in the hippocampus. D) Representative images from a double stain of Trem2 (red) and Clec7a (green). Percentage of area covered by Trem2 immunoreactivity (E) and ApoE immunoreactivity in Clec7a positive cells (F) was quantified in the hippocampus. G) Representative images of homeostatic microglia immunostained with P2ry12 in the hippocampus (Scale bar 50 µm). Total P2ry12 immunoreactivity area was analyzed using Imaris (H). Scale bar 30 µm. Data expressed as mean ± SD. One-way ANOVA with Tukey’s post hoc test (two-sided) was used for all statistical analysis *p<0.05, **p<0.01, p<0.001, ****p<0.0001.

**Figure S4.**
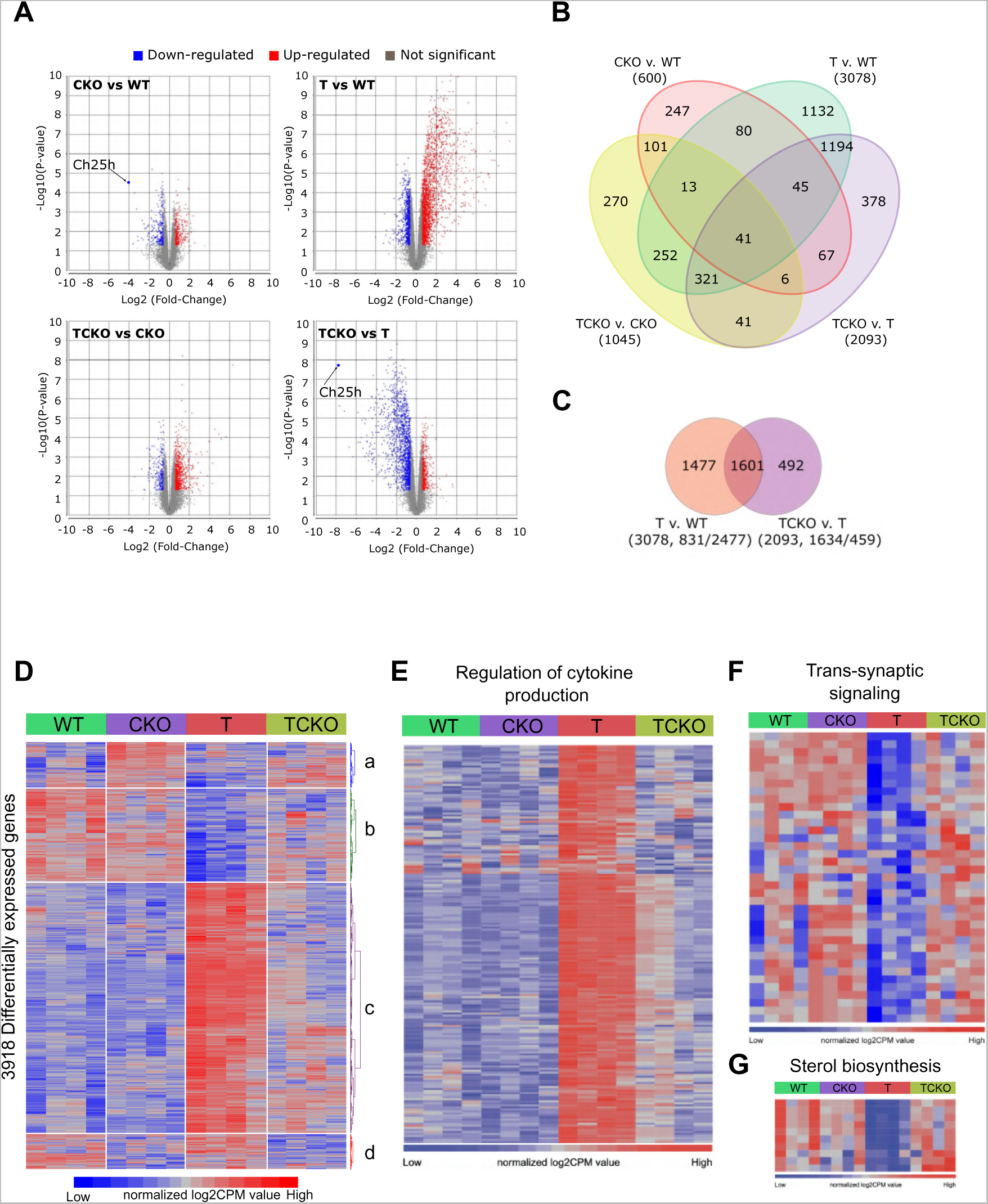
Transcriptomic changes induced by Ch25h deficiency in PS19 mice. Volcano plots of significant (p<0.05, Fold change (FC)>1.5) differentially expressed genes (DEGs) in the hippocampus of the comparisons CKO vs. WT, T vs. WT, TCKO vs. CKO, and TCKO vs. T mice (A). Venn diagram (B) showing DEGs in the comparisons CKO vs. WT, T vs. WT, TCKO vs. CKO, and TCKO vs. T mice. Venn diagram (C) showing DEGs in the comparisons T vs. WT and TCKO vs. T. (D) Heatmap of 3918 DEGs (p<0.05, FC>1.5) in WT, CKO, T, and TCKO mice (n=4 samples per group) showing the effects of Ch25h-deficiency and tau transgene. Cluster ‘a’ and ‘d’ show DEGs upregulated and downregulated in Ch25h-deficient (CKO and TCKO) groups, respectively. Cluster ‘b’ shows DEGs downregulated in T and restored in TCKO. Cluster ‘c’ shows DEGs upregulated in T and restored in TCKO. Heatmap of significant DEGs (FC>1.5, p<0.05) related to the GO terms ‘regulation of cytokine production’ (E), ‘trans-synaptic signaling’ (F) and ‘sterol biosynthetic process’ (G).

**Figure S5.**
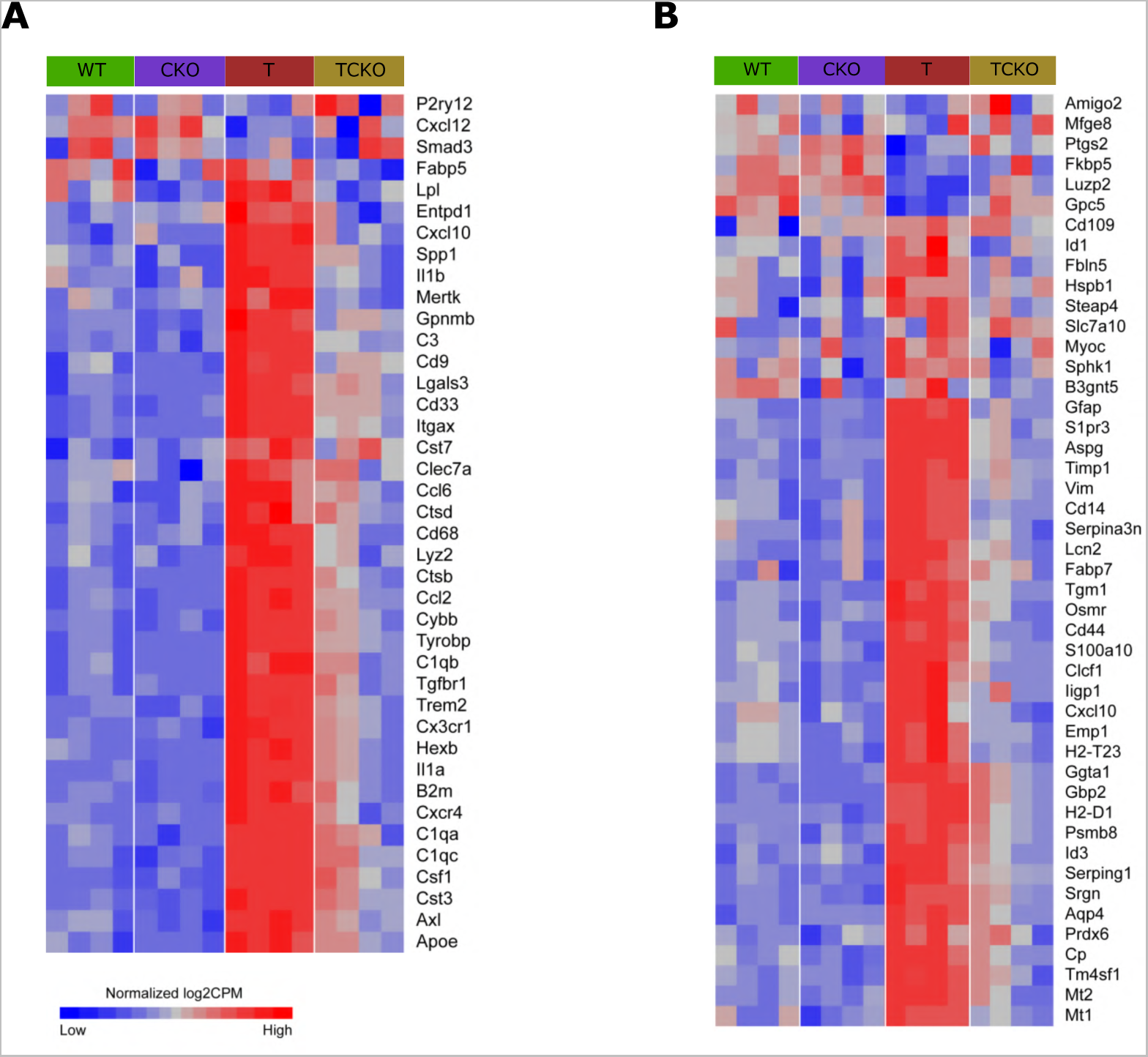
Ch25h deficiency in PS19 mice induces changes in genes of DAM and reactive astrocyte markers. Heatmap of significant DEGs (FC>1.5, p<0.05) of microglia (A) and astrocyte (B) markers.

**Figure S6.**
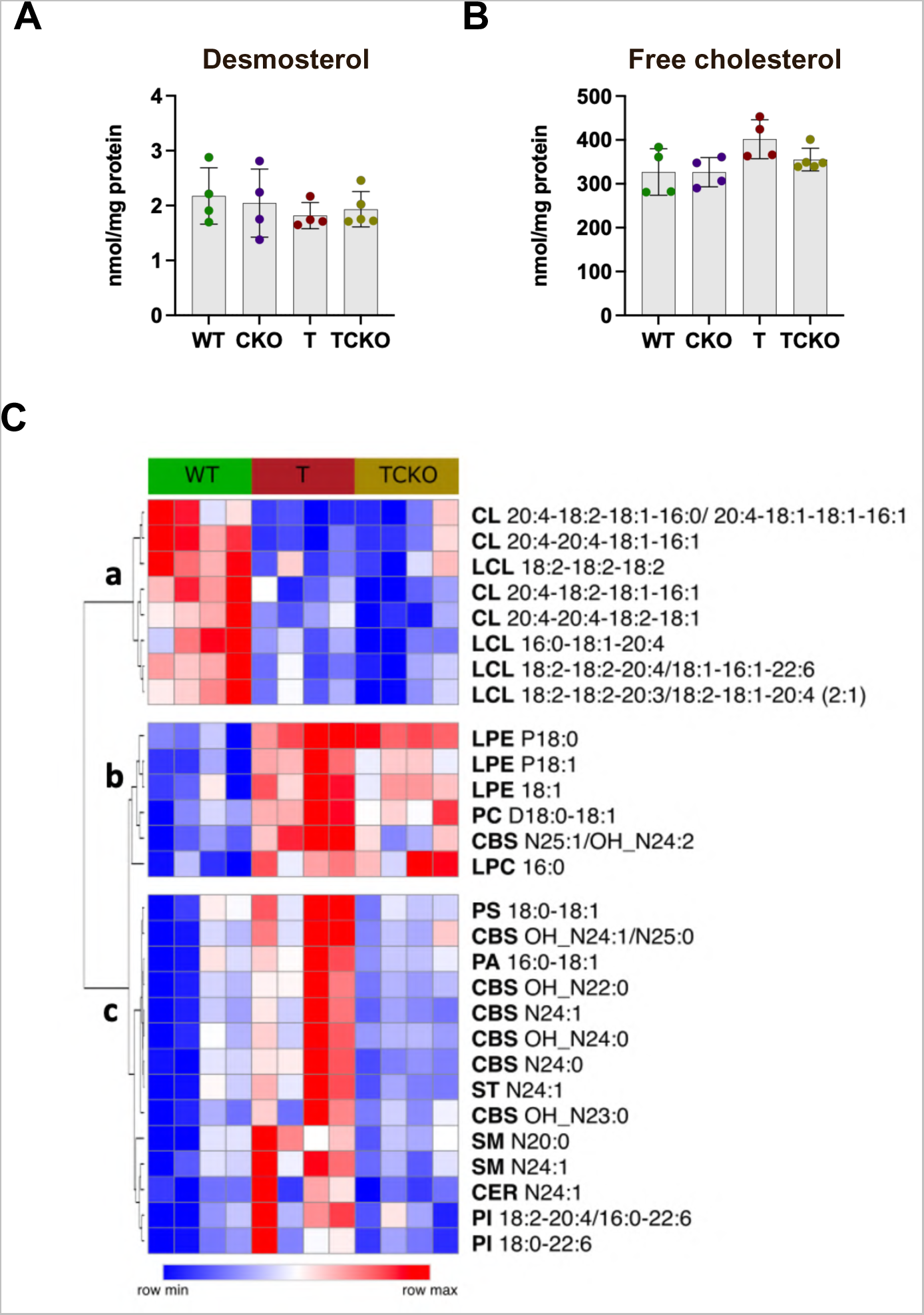
Ch25h deficiency in PS19 mice modulates levels of specific lipids without affecting free cholesterol levels. Quantification of demosterol (A) and free cholesterol (B) levels in the cortex of 9.5-month old WT, CKO, T and TCKO mice (n=4-5 samples per group). Lipid species were quantified (Supplementary Table VI) in the cortex of WT, T and TCKO mice (n=4 per group). One-way ANOVA with Tukey’s post hoc test (two-sided) was used for all statistical analysis. Heatmap of lipid species (C) that showed significant differences (p<0.05) between the study groups (row min-max). Clusters ‘a’ and ‘b’ show lipids reduced and increased in mice carrying the tau-transgene, respectively. Cluster ‘c’ shows lipids upregulated in T but restored in TCKO.

## SUPPLEMENTARY TABLE LEGENDS

**Supplementary Table I: Demographic information of the human brain tissue samples**

**Supplementary Table II: Total number of differentially expressed genes.** Comparisons are T vs WT, CKO vs WT, TCKO vs T and TCKO vs CKO.

**Supplementary Table III: List of differentially expressed genes.** Comparisons are T vs WT, CKO vs WT, TCKO vs T and TCKO vs CKO. Significant p-values are highlighted in yellow. Upregulated genes are in red and downregulated genes in green. Cluster IDs are as noted in Figure S4

**Supplementary Table IV: List of differentially expressed genes showing only the significant DEGs for each comparison (p-value <0.05 and fold change of 1.5).** Comparisons are T vs WT, CKO vs WT, TCKO vs T and TCKO vs CKO.

**Supplementary Table V: GO term analysis for up and down regulated genes in T vs WT and TCKO vs T comparisons.**

**Supplementary Table VI: Determination of lipid species in the cortex of WT, T and TCKO mice by LC/MS.**

**Supplementary Table VII: Summary of the data size of snRNA-seq results.**

**Supplementary Table VIII: Integrated average gene expression of cell-type specific markers.**

**Supplementary Table IX: Differential gene expression in microglia cluster. Supplementary Table X: Relative frequency of microglia cluster.**

## Notes

### Competing Interest Statement

SMP is a shareholder of Sage Therapeutics, Voyager Therapeutics, Karuna Therapeutics and Alnylam Pharmaceuticals and a Venture Partner at Third Rock Ventures. D.M.H. co-founded, has equity, and is on the scientific advisory board of C2N Diagnostics. D.M.H. is on the scientific advisory board of Denali, Cajal Neuroscience, and Genentech and consults for Asteroid Therapeutics.

