## Supplementary figures and images for "Microglial 25-hydroxycholesterol mediates neuroinflammation and neurodegeneration in a tauopathy mouse model"

### Supplementary

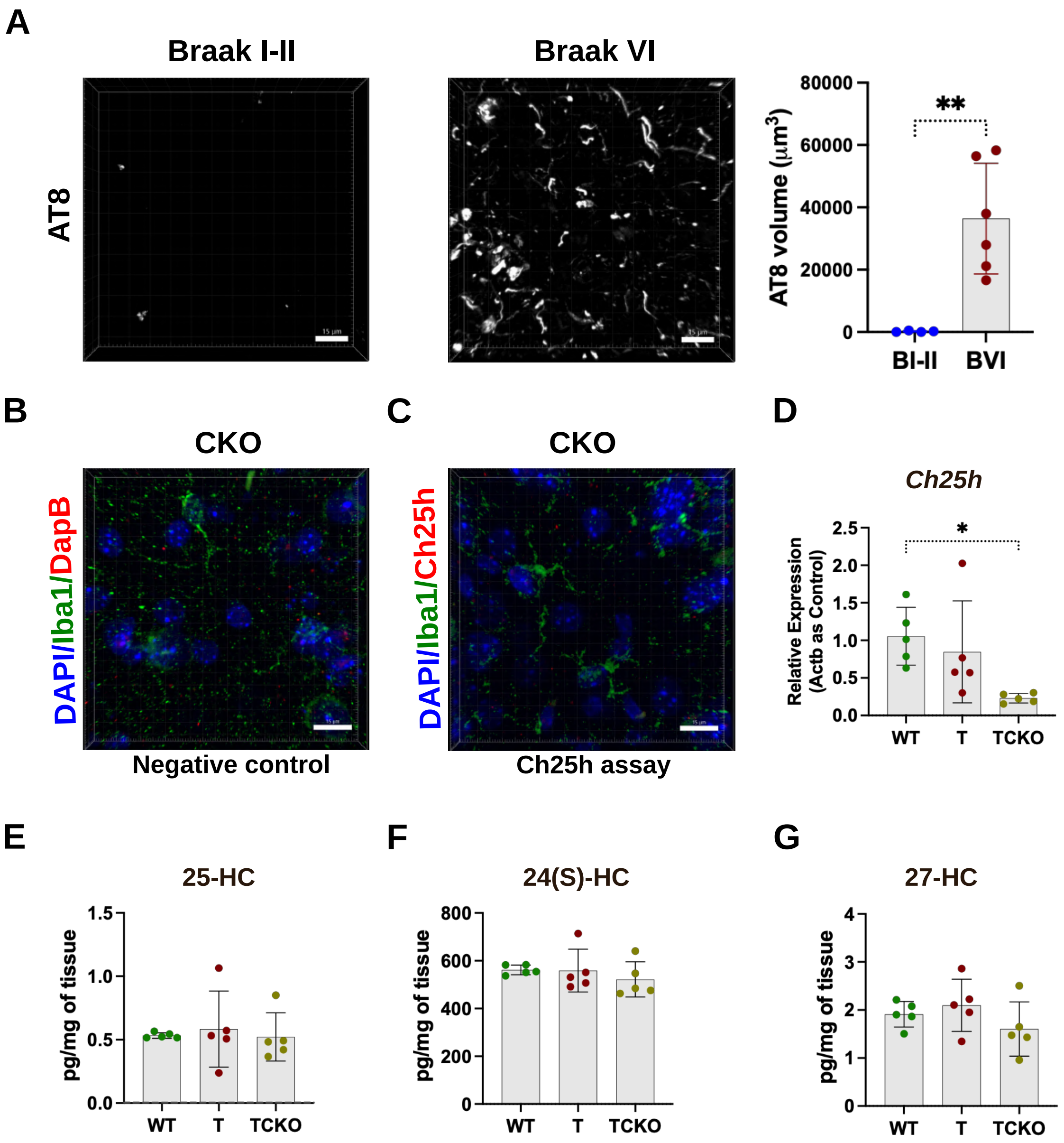

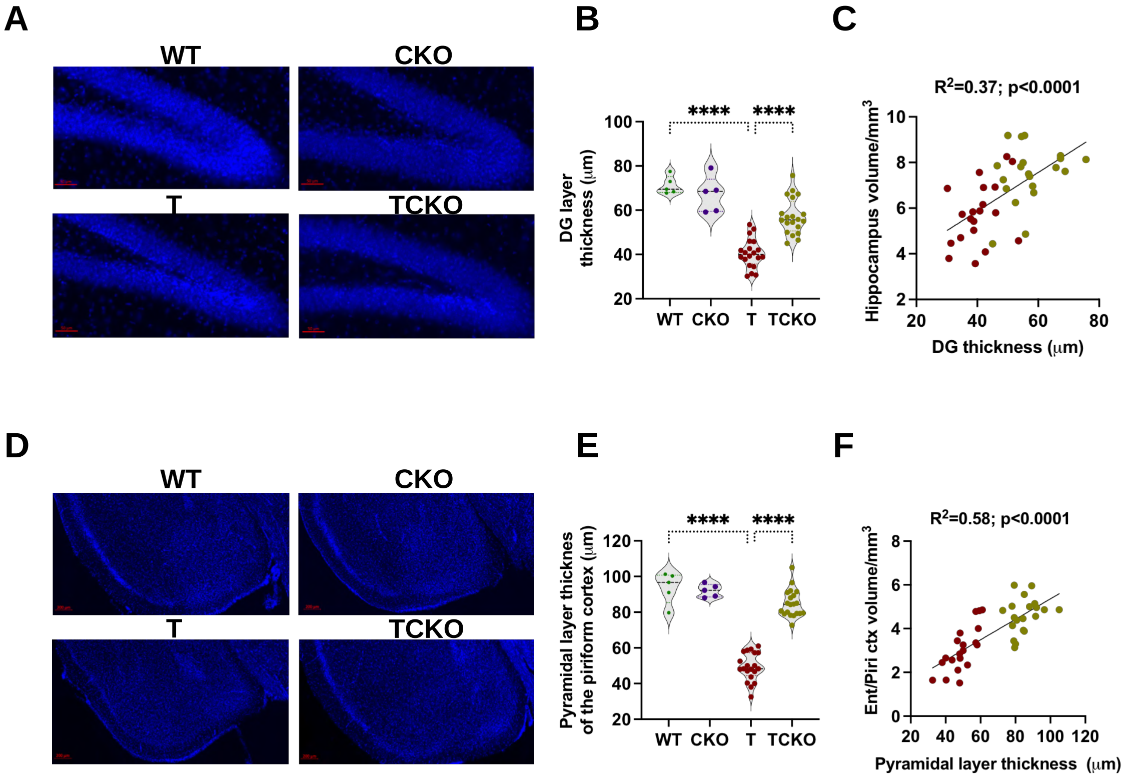

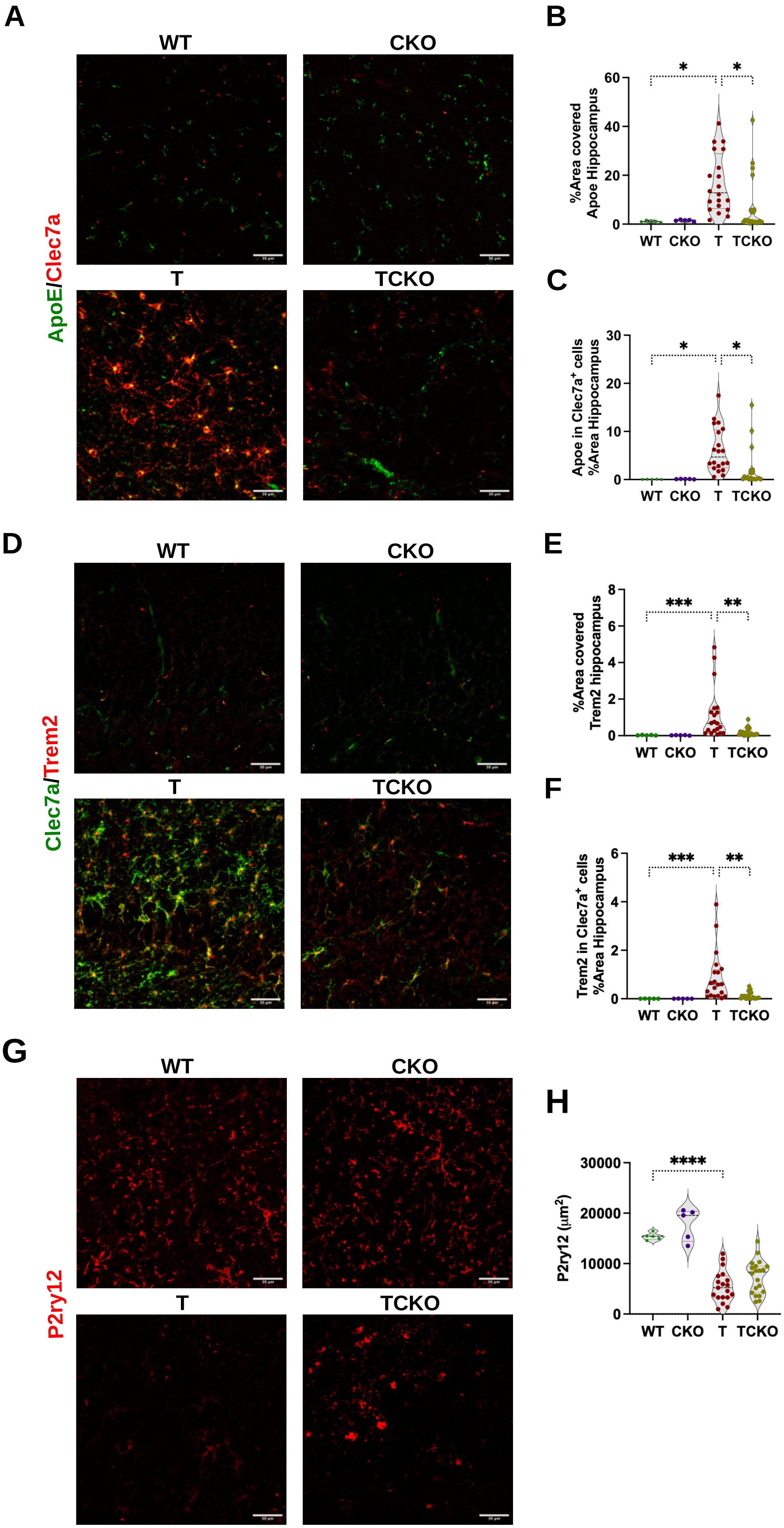

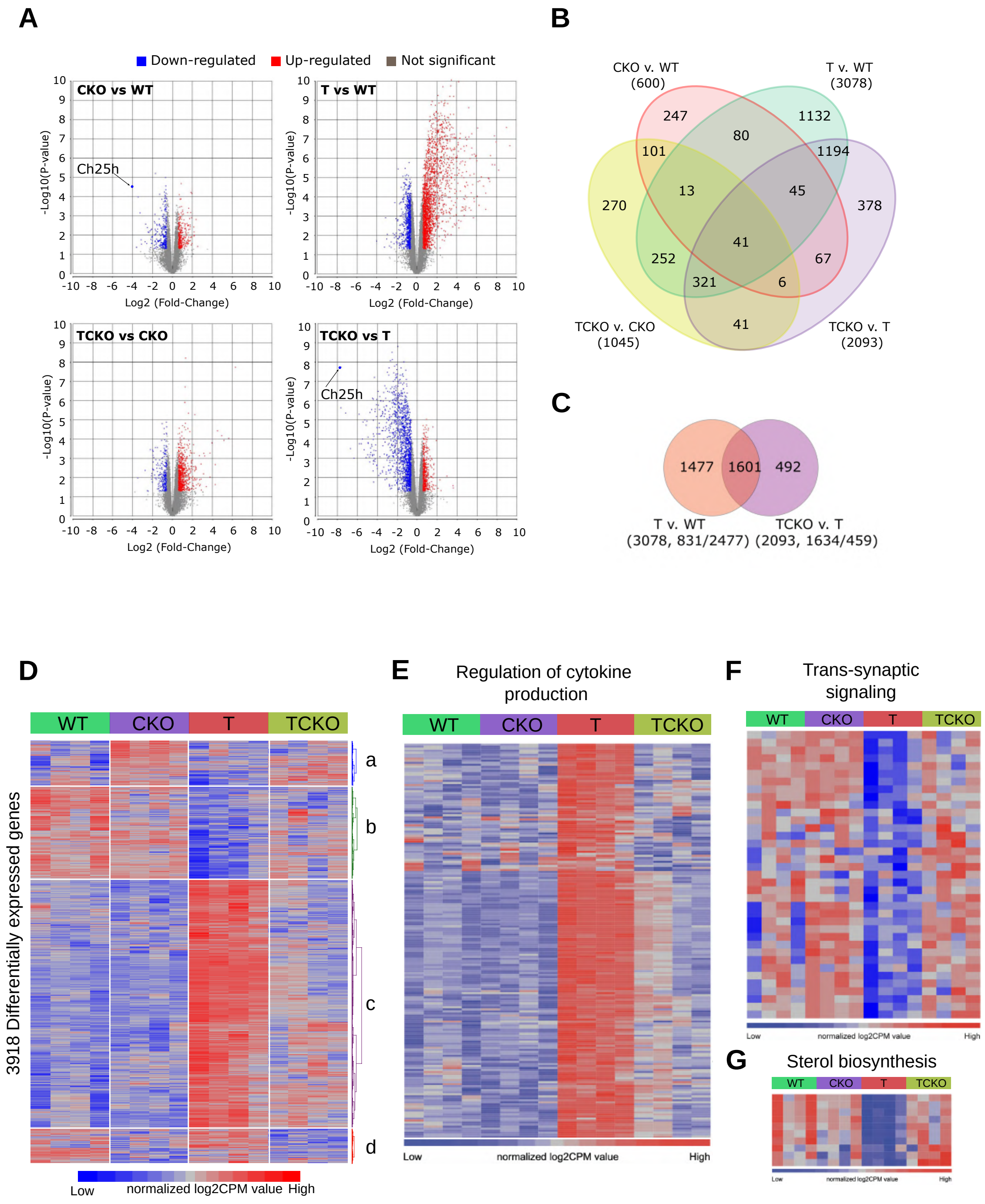

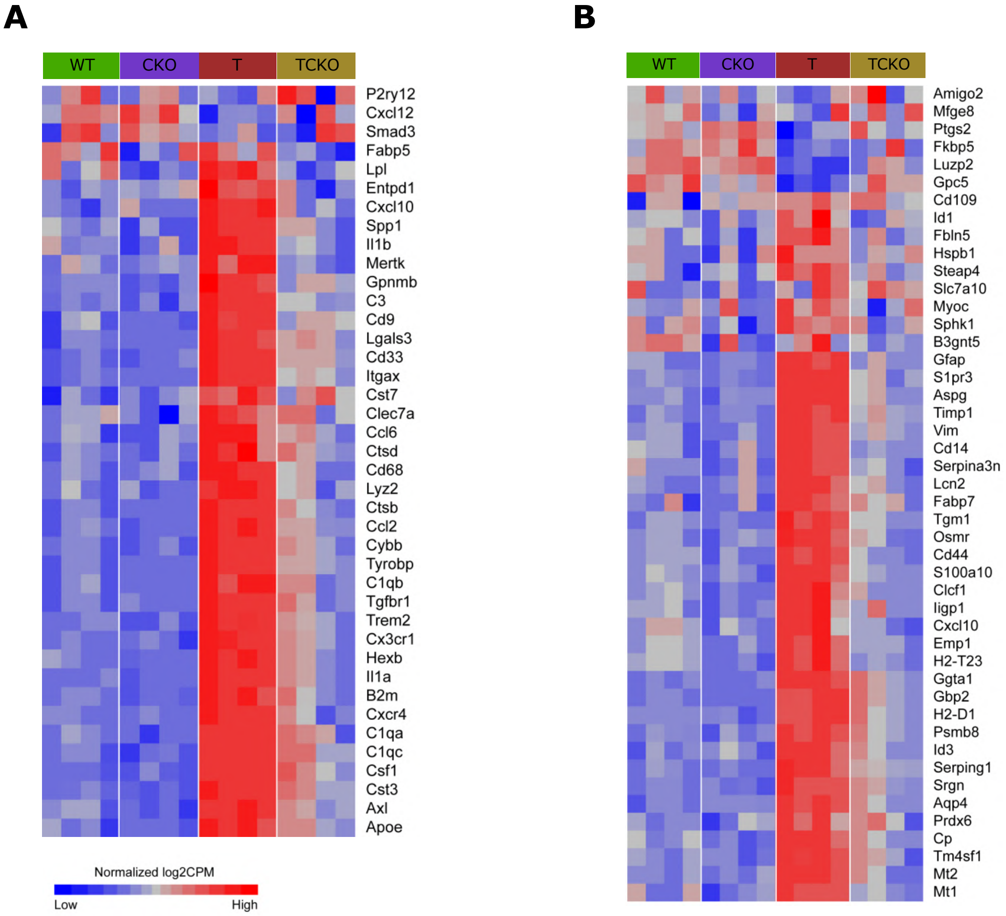

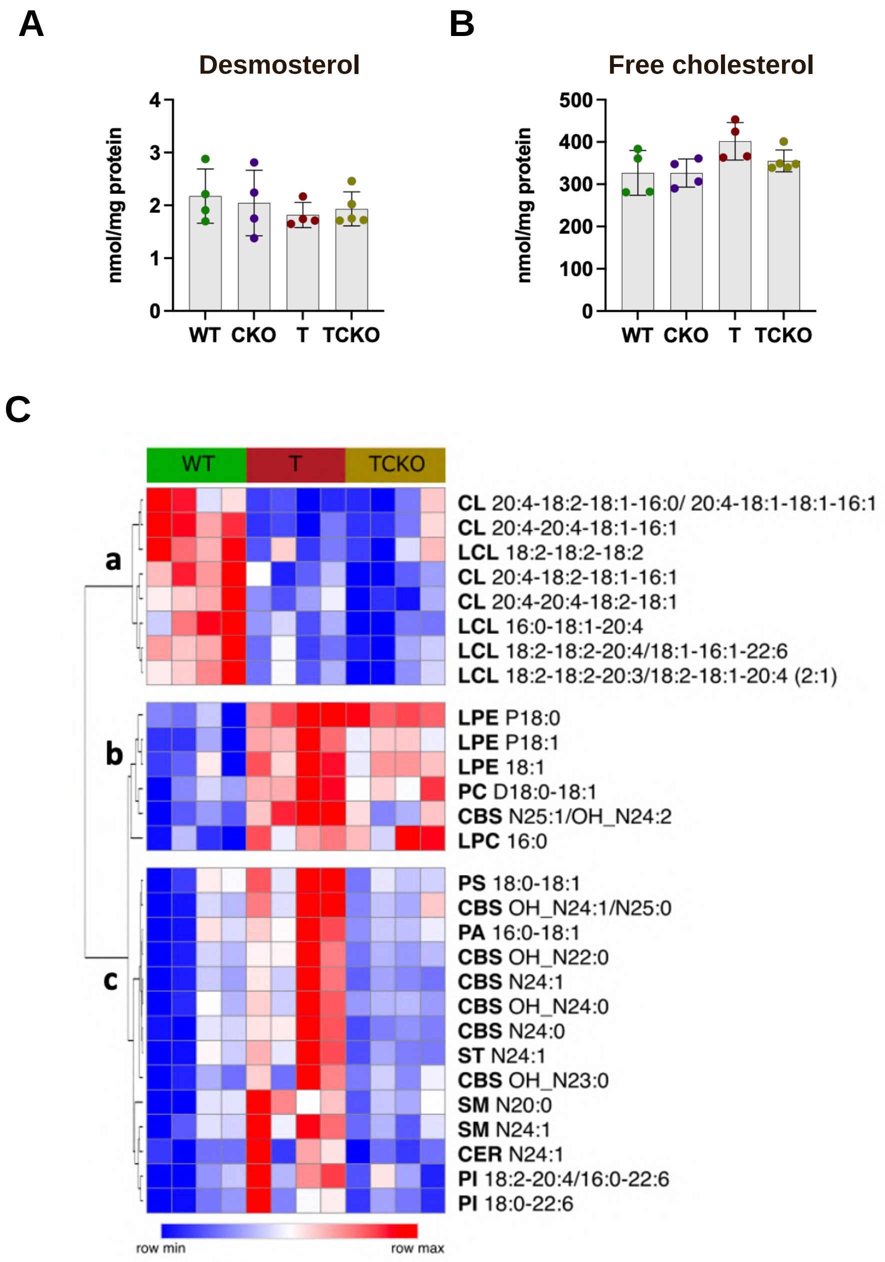
